# Catestatin ameliorates tauopathy and amyloidogenesis via adrenergic inhibition

**DOI:** 10.64898/2026.01.04.697519

**Authors:** Suborno Jati, Satadeepa Kal, Daniel Munoz-Mayorga, Kechun Tang, Debashis Sahoo, Xu Chen, Sushil K. Mahata

**Affiliations:** Department of Neuroscience, University of California San Diego, La Jolla, CA; Veterans Medical Research Foundation, San Diego, CA; Department of Medicine, University of California San Diego, La Jolla, CA; Department of Pediatrics, University of California San Diego, La Jolla, CA; Department of Computer Science and Engineering, University of California San Diego, La Jolla, CA; VA San Diego Healthcare System, San Diego, CA

**Author notes:** Co-correspondence: Sushil K. Mahata Xu Chen Suborno Jati. Lead Contact: Sushil K. Mahata.

## Abstract

Neurodegenerative disorders like Alzheimer’s disease (AD), Corticobasal Degeneration (CBD), and Progressive Supranuclear Palsy (PSP) are characterized by Tau aggregation, synaptic dysfunction, neuroinflammation, and progressive cognitive decline. Although metabolic dysregulation and neuropeptide imbalance have been linked to these disorders, the functional consequences of such imbalance and its potential for therapeutic reversal remain poorly understood. Our previous work identified chromogranin A (CgA), which encodes a pro-hormone for several metabolic peptides, as a key regulator of Tau pathology. Here, we investigate Catestatin (CST), a CgA-derived peptide that is a potent inhibitor of catecholamine release and has been shown to increase insulin sensitivity and lower peripheral blood pressure. We report significant reductions in CST levels in the hippocampus and cortex of AD brains, as well as in the frontal cortex of CBD and the basal ganglia of PSP. Supplementing CST in cortical neuronal cultures and organotypic slice cultures (OTSC) decreased Tau phosphorylation and aggregation. *In vivo*, CST administration in PS19 Tauopathy mice reduced pathological Tau species, attenuated gliosis, and improved cognitive function. CST treatment also lowered amyloid plaque burden and neuroinflammation in 5xFAD mice. Mechanistically, CST decreased epinephrine (EPI) levels in both PS19 and 5xFAD mice and suppressed downstream protein kinase A (PKA) hyperactivation in PS19 and OTSC. These findings reveal a previously unrecognized neuropeptidergic mechanism linking CST deficiency to elevated adrenergic receptor (ADR)-EPI–PKA stress signaling and Tauopathy-driven neurodegeneration, suggesting CST replacement as a promising therapeutic approach.

## Introduction

Tauopathies represent a diverse class of neurodegenerative disorders defined by abnormal post-translational modification ^1^, misfolding ^2^, and aggregation of the microtubule-associated protein Tau ^3^. Alzheimer’s disease (AD), the most prevalent Tauopathy, is also characterized by extracellular amyloid-β (Aβ) plaques and progressive cognitive impairment and brain atrophy ^4^. Other primary Tauopathies, such as corticobasal degeneration (CBD) and progressive supranuclear palsy (PSP), display region-specific accumulation of pathological Tau isoforms and share overlapping clinical features, including motor dysfunction, executive deficits, and neuronal loss in affected cortical and subcortical regions ^5^. Although extensive research has elucidated upstream triggers of Tau dysfunction - including aberrant kinase activation, neuroinflammation, and impaired proteostasis - the mechanisms linking systemic metabolic signals to Tau pathology remain poorly understood.

Chromogranin A (CgA), an acidic secretory pro-protein stored in neuroendocrine dense-core secretory vesicles (DCVs) with resident hormones ^6,7^, is increasingly recognized as a modulator of neurodegenerative processes ^8^. CgA levels are elevated in AD brain and cerebrospinal fluid (CSF) ^8–10^, and proteolytic processing of CgA generates a class of bioactive peptides with diverse effects on inflammation, metabolism, and synaptic function. These include Catestatin (CST: hCgA352-372), a potent inhibitor of catecholamine release ^11,12^; Pancreastatin (PST: hCgA250-301), a metabolic antagonist of insulin ^13–15^; and Vasostatin (hCgA1-76), a vasodilator peptide ^16^. Dysregulation of CgA-derived peptides has been reported in cardiometabolic disease ^17–19^, immune regulation ^15,20^ and Alzheimer’s disease ^21^, however, their mechanistic role in neurodegeneration - particularly in Tauopathies - remains largely unexplored.

Therapeutic strategies for AD and related Tauopathies have advanced substantially in recent years. Small-molecule inhibitors targeting amyloid production (BACE1 inhibitors) or Tau kinases (GSK3β, CDK5 inhibitors) have shown promising preclinical efficacy but have had limited clinical translation due to toxicity or a lack of cognitive benefit ^22–26^. Immunotherapies, including anti-Aβ monoclonal antibodies such as aducanumab, lecanemab, and donanemab, can reduce amyloid burden yet provide modest and variable clinical improvement ^27,28^. Anti-Tau antibodies and vaccines are under active investigation but have yet to demonstrate consistent disease-modifying impact ^29^. In parallel, peptide-based therapies, which offer high target specificity and favorable safety profiles, are gaining traction in neurodegeneration. Peptides such as NAP (davunetide), selenopeptides, bradykinin analogs, and mitochondrial-protective peptides (SS-31) have shown neuroprotective effects in preclinical studies ^30–33^. Despite these advances and the known neuropeptide imbalance in the neurodegenerative brain ^34,35^, no peptide therapy has yet been established to directly modulate Tau pathology through endogenous neuroendocrine pathways.

In this study, we investigated whether CST with established anti-adrenergic ^11,12^, anti-inflammatory ^20,36^, and metabolic regulatory functions ^20,37^ plays a protective role in Tauopathy and amyloid pathology. We first quantified CST levels in human AD, CBD, and PSP brain tissue, revealing marked reductions across disease-affected regions. We then assessed the therapeutic potential of CST using complementary systems: cortical neuronal cultures, hippocampal organotypic slice cultures, PS19 Tauopathy mice, and 5xFAD amyloid mice. Across these models, CST supplementation mitigated Tau phosphorylation and aggregation, reduced amyloid burden, suppressed neuroinflammation, restored hippocampal volumes, normalized cell-type composition in snRNA-seq, and improved cognitive and motor functions. Finally, we examined whether CST modulates epinephrine (EPI)-driven protein kinase A (PKA) activation, a pathway known to promote Tau phosphorylation, and demonstrated that CST reduces EPI levels and corresponding PKA signaling *in vivo* and *ex vivo*.

Together, these findings identify CST as a previously unrecognized regulator of Tau and amyloid pathology and provide a strong mechanistic rationale for CST-based peptide therapy in AD and related neurodegenerative disorders.

## Results

### CST levels decline across AD, CBD, and PSP

To determine whether CST dysregulation associates with human neurodegeneration, we quantified CST levels in postmortem AD cortex and hippocampus using a commercially available kit ^38^. Quantitative ELISA for CST in the pre-frontal cortex (PFC) lysates revealed a significant decrease in CST level in Braak stage 6 patients (patient details in **S-Table. 1**) compared to Braak stages 0-2 (**Fig. 1A**). Lower CST abundance correlated with worse cognitive performance: CST levels in PFC showed positive association with Mini-Mental State Examination (MMSE) scores (PFC: R² = 0.3357, p = 0.0008; **Fig. 1B**). A similar reduction of CST level was observed in hippocampus/entorhinal cortex (HC/EC) samples (**Fig. 1C**) and lowered CST level in HC/EC also correlated with lower MMSE scores (HC/EC: R² = 0.2526, p = 0.0047; **Fig. 1D**). Reduced CST level in AD brains was further corroborated by immunohistochemistry (IHC) with CST showing significant reduction of CST in the cell body of Dentate Gyrus of Braak 6 stage patient compared to Braak 0-2 (**Fig. 1E&F**). As expected, AD PFC samples with low CST also exhibited increased pTau (S202/T205) levels by immunostaining (**S-Fig. 1A**), linking CST deficiency to the neuropathological severity of late-stage AD. These findings suggest that CST, as a regulatory peptide, is negatively correlated with AD pathophysiology.

**Figure 1.**
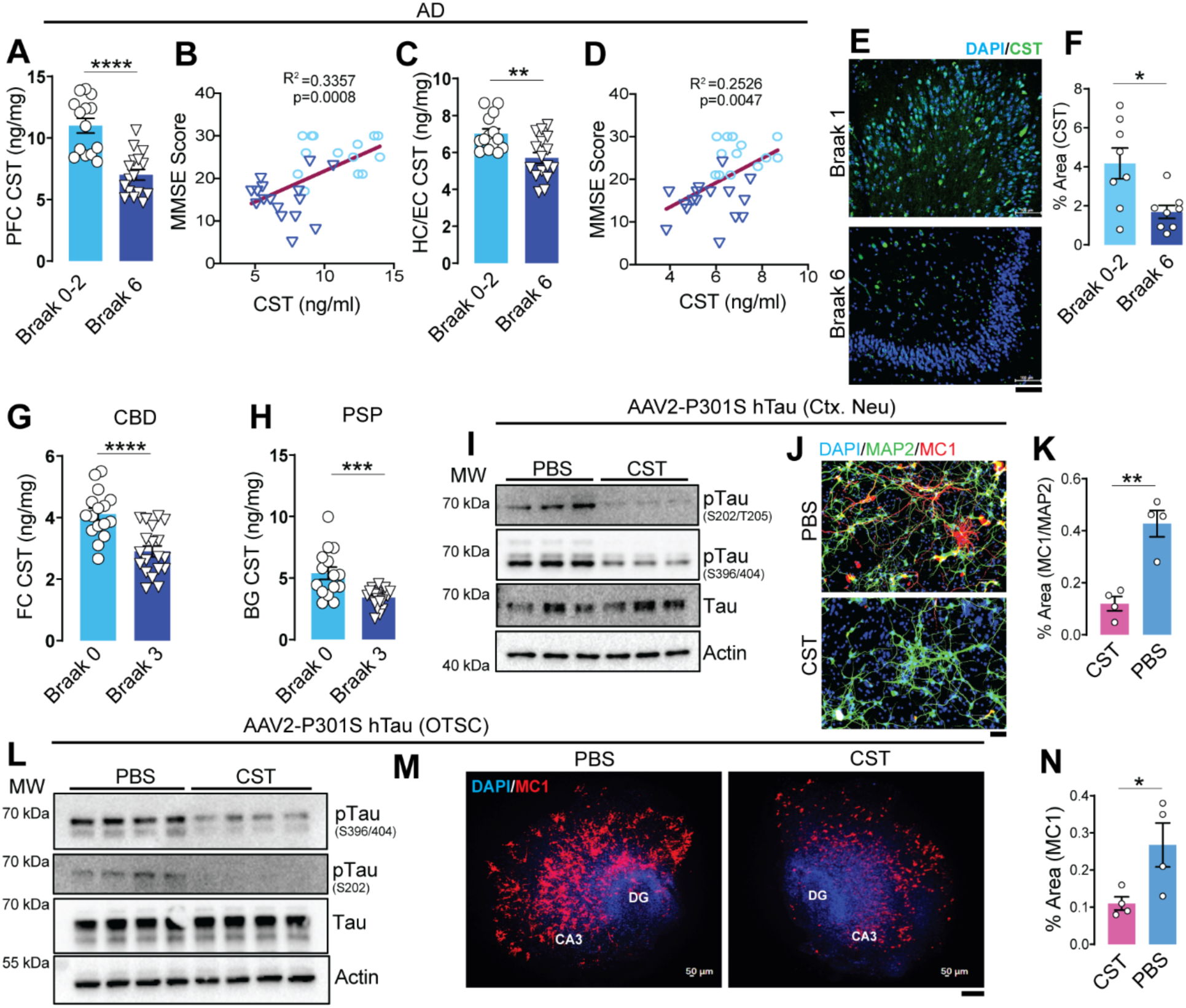
CST levels are reduced across Tauopathies, and AD-spectrum disorders, and CST supplementation suppresses pathological Tau accumulation. CST quantification in human Alzheimer’s disease (AD) cortex shows significantly reduced CST in the (A) prefrontal cortex (PFC) and (C) hippocampus/entorhinal cortex (HC/EC) of Braak VI (n=16) vs. Braak 0–II (n=14) subjects. CST levels positively correlate with Mini-Mental State Examination (MMSE) scores in (B) PFC and (D) HC/EC, linking CST deficiency to cognitive decline. (E&F) Immunostaining of CST in postmortem tissue reveals reduced CST level in Braak VI (n=8) hippocampus compared with Braak 0–II (n=8); quantification shown in (**F**). Scale bar = 100µm **(G&H)** CST levels are also significantly reduced in cortical tissue from (**G**) corticobasal degeneration (CBD; Braak 0: n=16, Braak 3: n=19) and (**H**) basal ganglia from progressive supranuclear palsy (PSP; Braak 0: n=15, Braak 3: n=19), indicating CST loss as a shared feature of 4R-Tauopathies. **(I)** In cortical neuron cultures transduced with AAV2–P301S hTau, CST treatment decreases hyperphosphorylated Tau species (S202/T205 and S396/S404) without altering total Tau (n=3). **(J–K)** Immunocytochemistry confirms CST-mediated suppression of MC1+ misfolded Tau in P301S hTau–expressing neurons; quantified as MC1/MAP2+ area in (n=4). Scale bar = 50µm. **(L–N)** In organotypic hippocampal slice cultures (OTSC) expressing AAV2–P301S hTau (n=4), (**L**) CST reduces pTau (S202) and pTau (S396/S404) accumulation and (**M&N**) diminishes MC1+ Tau pathology in DG and CA3 regions. Scale bar = 200µm *Data are mean ± SEM; statistical significance indicated as *p < 0.05, **p < 0.01, ***p < 0.001, ***p < 0.0001.

CST is one of the several bioactive peptides derived from Chromogranin A (CgA). Out of these peptides, CST and PST are known to have antagonistic metabolic functions ^13^. We therefore evaluated PST abundance in AD samples. PST levels were increased in PFC and HC/EC of Braak stage 6 cortex compared to Braak 0–2 (**S-Fig. 1B-C**), suggesting a CST↓ / PST↑ imbalance in advanced AD.

We next examined whether CST dysregulation extends to primary Tauopathies. In the CBD frontal cortex (FC), CST levels were significantly decreased at Braak stage 3 (Patient details in **S-Table. 2**) compared with Braak 0 (**Fig. 1G**). In contrast, PST levels were increased (**S-Fig. 1D**). Similarly, in PSP, CST levels in the basal ganglia (BG) were reduced at Braak stage 3 (patient details in **S-Table. 3**) versus Braak 0 (**Fig. 1H**), supporting a consistent shift in the balance of CgA-derived peptides across various Tauopathies.

### CST supplementation reduces pathological Tau accumulation *in vitro* and *ex vivo*

To directly test whether CST modulates Tauopathy, we first tested CST treatment on primary neurons transduced with AAV2-P301S human Tau and inoculated with preformed Tau fibrils (PFFs, K18/PL). CST treatment markedly reduced phosphorylation at pTau (S202/T205 and S396/404), as assessed by western blotting (**Fig. 1I, S-Fig. 1E&F**). CST also suppressed the accumulation of misfolded Tau species (MC1+), as seen by IHC (**Fig. 1J&K**). We further assessed the effect of CST treatment in hippocampal organotypic slice culture (OTSC) transduced with AAV2-P301S human Tau (hTau) and seeded with PFFs. Similar reductions in pTau (S202/T205, S396/404) and misfolded Tau aggregates (MC1+) were observed in CST-treated OTSCs compared to PBS controls (**Fig. 1L-N, S-Fig. 1G&H**). In contrast, PST treatment increased pTau levels (**S-Fig. 1I&J**). These *in vitro* and *ex vivo* data suggest that CST supplementation may reduce pathological Tau levels.

### CST treatment reduces hippocampal atrophy and rescues behavioral deficits in PS19 mice

We next evaluated whether CST supplementation can lower Tau phosphorylation and aggregation *in vivo* in the PS19 Tau transgenic mouse model. In the cortex and plasma of PS19 mice at 9 months of age, we observed reduced CST levels compared to non-transgenic (nTg) controls (**S-Fig. 2A&B**), reminiscent of levels observed in Tauopathy patients. We next investigated the effect of CST supplementation in PS19 mice. Mice were dosed with CST (0.5 µg/g body weight, i.p., 3x/week) beginning at 5 months of age- when early neuropathological changes emerge - and continued treatment until 9.5 months (**Fig. 2A**). Following intraperitoneal (i.p.) administration at 5 mg/kg in mice (**S-Fig. 2D)** CST showed rapid systemic absorption but diverged markedly in their plasma and brain distribution profiles (**S-Fig. 2E&F**). CST achieved a plasma *C*max of 148.3 ng/ml at 0.25 h and displayed a relatively short terminal half-life (t1/2 = 1.28 h). Notably, CST penetrated the CNS efficiently, with brain concentrations peaking at 44.5 ng/g and brain AUC exceeding plasma AUC (**brain/plasma AUC ratio, Kp = 1.694, S-Table. 4**). At the endpoint (9.5 months), saline-treated PS19 mice displayed pronounced hippocampal atrophy. In contrast, CST-treated PS19 mice exhibited significantly preserved hippocampal structure, with volumes indistinguishable from those of nTg controls (**Fig. 2B&C**). Morphometric examination of hippocampus revealed the following: decreased thickness of dentate gyrus (DG) granule cell layer and CA1 pyramidal cell layer by ∼25% and ∼37% (**S-Fig. 2C**), respectively, in PS19 mice; increased thickness of both the DG and CA1 layers by ∼42% and ∼46%), respectively, following CST treatment, indicating a robust protection against neurodegeneration.

**Figure 2.**
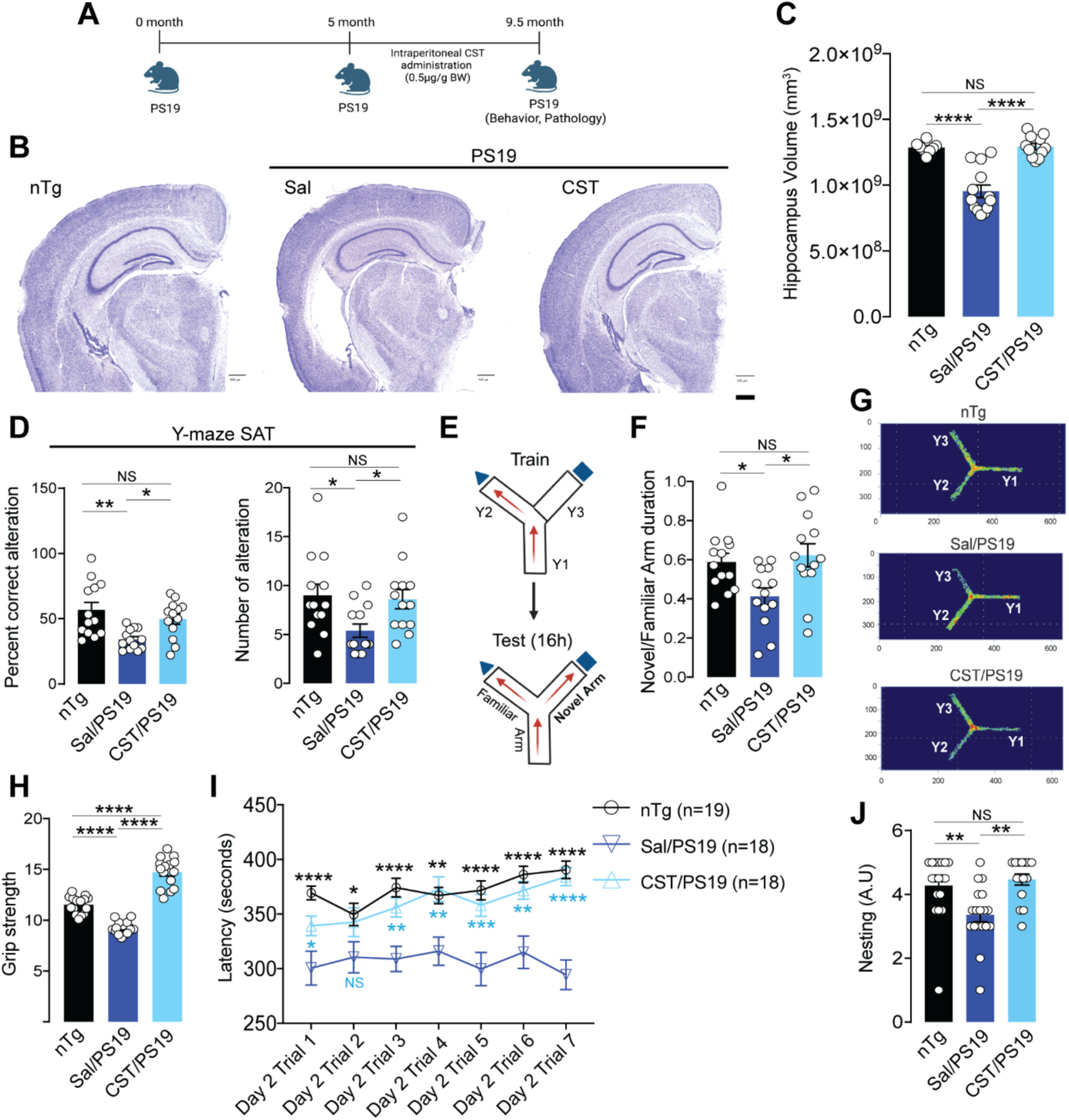
CST treatment rescues hippocampal atrophy and improves cognitive, motor, and innate behaviors in PS19 mice. **(A)** Experimental design showing intraperitoneal CST administration (0.5 µg/g BW) to PS19 mice from 5 to 9.5 months of age, followed by behavioral and pathological assessments. **(B&C)** CST preserves hippocampal structure in PS19 mice. (**B**) Representative Nissl-stained coronal sections and (**C**) volumetric analyses in nTg (n=8), Sal/PS19 (n=13), and CST/PS19 (n=11) reveal marked protection against hippocampal atrophy in CST-treated PS19 mice compared with saline-treated PS19 controls. **(D–F)** CST improves hippocampus-dependent working and spatial memory. In the Y-maze spontaneous alternation test (SAT), CST-treated PS19 mice exhibited (**D**) increased percent alternation and number of alterations (n=13). (**E**) In the two-trial Y-maze test, (**F**) CST-treated mice show greater novel-arm exploration than saline-treated PS19 mice (n=13). **(G)** Representative heatmaps of arm exploration illustrate superior spatial memory in CST-treated PS19 mice. **(H–J)** CST ameliorates motor deficits and restores innate behaviors. CST-treated PS19 mice show (**H**) improved grip strength (n=15), (**I**) significantly enhanced rotarod performance across training trials (nTg: n=19, Sal/PS19: n=18, CST/PS19: n=15), and (**J**) improved nest-building behavior compared with saline-treated PS19 animals (nTg: n=18, Sal/PS19: n=18, CST/PS19: n=15). *Data represent mean ± SEM. Significance levels: *p < 0.05, **p < 0.01, *\*\*\**p < 0.001, ****p < 0.0001; NS, not significant

Because PS19 mice begin to exhibit cognitive and motor deficits between 7–8 months, we performed behavioral tests at 7 months to assess whether early CST intervention improves functional outcomes. In the Y-maze Spontaneous Alternation Test (SAT), saline-treated PS19 mice showed a marked reduction in percent alternation relative to nTg controls. CST treatment significantly increased alternation performance, restoring it to nTg levels (**Fig. 2D**). The number of arm entries, a measure of exploratory activity, also showed improvement with CST treatment, indicating preserved working memory and spatial strategy.

To further assess working memory, we used a two-trial Y-maze paradigm (**Fig. 2E**). CST-treated PS19 mice demonstrated a significantly higher preference for the novel arm compared to saline-treated PS19 animals, reflecting enhanced short-term spatial memory (**Fig. 2F**). Heatmaps of arm exploration showed that CST-treated PS19 mice preferentially explored the novel arm, similar to nTg controls (**Fig. 2G**).

CST treatment also improved motor and innate behaviors. The forelimb grip strength test showed that CST-treated PS19 mice had substantially higher grip strength than saline-treated PS19 mice (**Fig. 2H**). In the multi-trial rotarod test, CST-treated PS19 mice exhibited significantly longer latency to fall across all trials, performing comparably to nTg controls (**Fig. 2I**). In addition, in the nesting assay, CST-treated PS19 mice showed higher nesting scores than saline-treated mice, indicating enhanced goal-directed and home-cage behavior (**Fig. 2J**).

Together, these findings demonstrate that CST supplementation ameliorated hippocampal atrophy and improved cognitive behaviors and motor function in PS19 mice. These data strongly support CST as a therapeutic agent for treating Tauopathy.

### CST treatment reduces Tau phosphorylation and aggregation in PS19 mice

After CST treatment in PS19, we analyzed Tau phosphorylation and aggregation using immunoblotting and IHC. Immunoblotting of cortical lysates showed that CST treatment markedly reduced levels of pTau at S202/T205 and S396/404 compared to saline-treated PS19 mice **(Fig. 3A-C**). A similar reduction in Tau phosphorylation was observed in the hippocampus (**Fig. 3D–F**).

**Figure 3.**
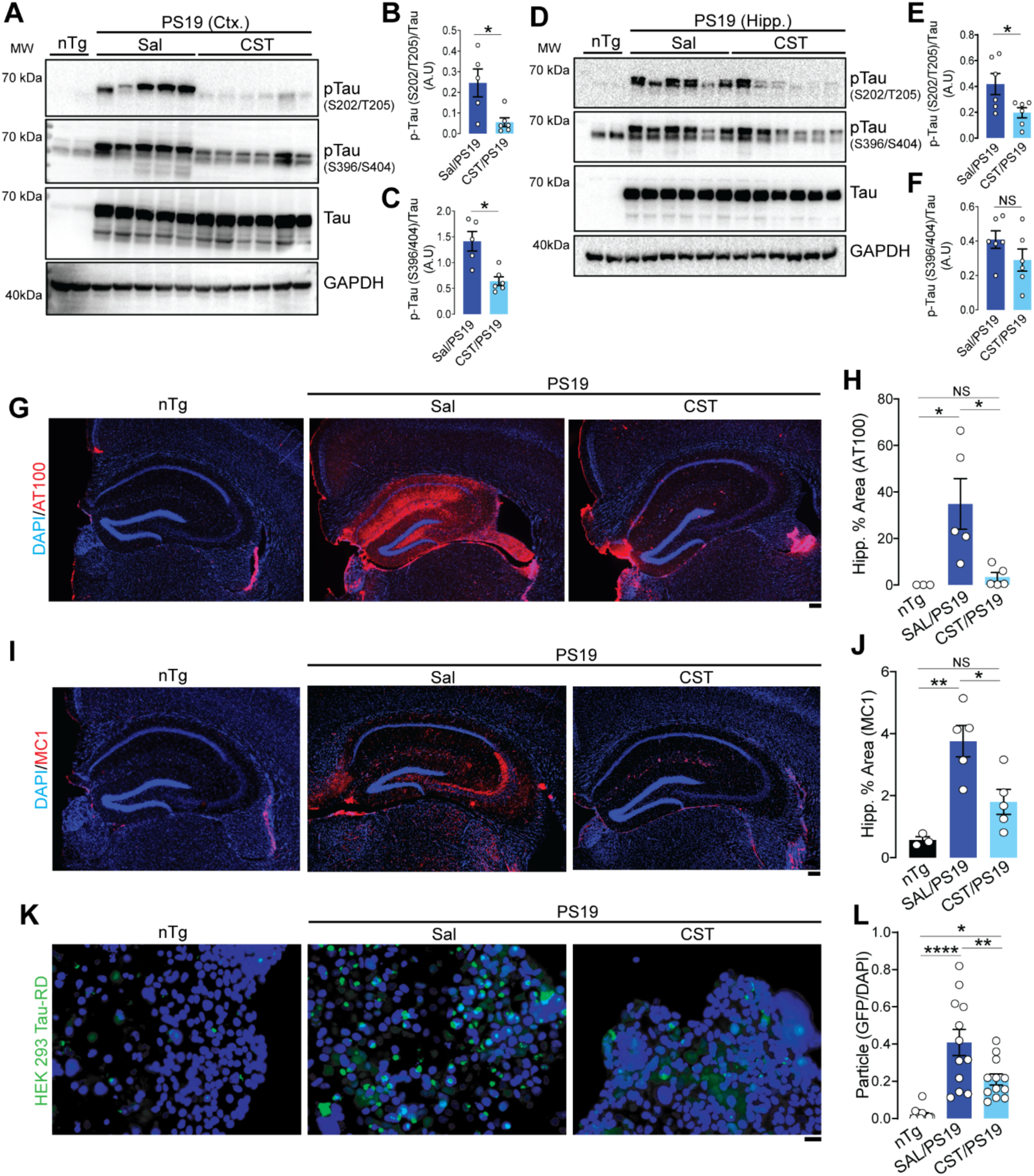
CST reduces Tau phosphorylation, misfolding, and seeding competence in PS19 mice. (A–C) Immunoblot analysis of cortical lysates shows elevated phospho-Tau species (S202/T205, S396/404) in saline-treated PS19 mice, both of which are significantly reduced following CST treatment (nTg: n=2, Sal/PS19: n=5, CST/PS19: n=6). **(D–F)** CST exerts similar biochemical effects in the hippocampus, where CST administration markedly decreases pathological Tau phosphorylation at S202/T205 and S396/S404 in PS19 mice (nTg: n=2, Sal/PS19: n=6, CST/PS19: n=6). (**G&H)** Hippocampal immunostaining with AT100 (recognizing pTau T212/S214) reveals substantial accumulation of pathological Tau in saline-treated PS19 mice; CST treatment significantly reduces the AT100-positive area, indicating reduced late-stage Tau pathology (nTg: n=3, Sal/PS19: n=5, CST/PS19: n=5). Scale bar = 200µm. (**I&J)** CST attenuates misfolded Tau burden. MC1 immunostaining demonstrates robust conformational Tau pathology in saline-treated PS19 mice, whereas CST markedly diminishes MC1-positive aggregates throughout the hippocampus (nTg: n=3, Sal/PS19: n=5, CST/PS19: n=5). Scale bar = 200µm. (**K&L)** CST suppresses Tau seeding competency. In HEK293 Tau-RD FRET biosensor assay, cortical lysates from saline-treated PS19 mice induce strong Tau aggregation. In contrast, lysates from CST-treated PS19 brains elicit significantly fewer GFP-positive aggregates, indicating reduced seeding activity (nTg: n=4, Sal/PS19: n=4, CST/PS19: n=4). Scale bar = 50µm. A.U.: Arbitrary Unit. Data represent mean ± SEM; significance indicated as *p < 0.05, **p < 0.01, *\*\*\**p < 0.001, ****p < 0.0001; NS, not significant.

To assess region-specific Tau pathology *in situ*, we examined hippocampal sections using the AT100 antibody, which recognizes the pTau T212/S214 associated with late-stage Tau pathology. Saline-treated PS19 mice showed extensive AT100 immunoreactivity throughout the hippocampus, especially in the CA1 region, whereas CST-treated mice displayed a substantial reduction in AT100-positive area (**Fig. 3G&H, S-Fig. 3A&B**). This decrease in AT100 staining corroborates the reduction in pTau shown by immunoblotting.

We next assessed misfolded Tau species using the conformation-specific MC1 antibody. Saline-treated PS19 mice showed prominent MC1 staining in the hippocampus - particularly in CA3 - whereas CST-treated PS19 mice exhibited significantly reduced MC1-positive area, comparable to nTg levels (**Fig. 3I&J, S-Fig. 3C&D**). These data demonstrate that CST reduces pathological Tau levels.

To measure the load of seed-competent Tau species in CST and saline-treated PS19, we employed a FRET-based HEK293T Tau-RD-CFP/YFP biosensor assay, which generates FRET signal when inoculated with Tau seeds. Cortical extracts from nTg mice produced no detectable seeding activity, as expected. Cortical extracts from saline-treated PS19 brains produced a robust FRET signal, indicative of high levels of aggregation-competent Tau seeds. In contrast, CST-treated PS19 brain extracts induced a markedly lower FRET response, reflecting a reduced abundance of transmissible Tau seeds **(Fig. 3K&L**).

Together, these biochemical, histological, and biosensor-based analyses demonstrate that CST treatment significantly suppresses Tau hyperphosphorylation, misfolding, and seeding competence in PS19 mice. These findings strongly demonstrate that CST treatment reduced pathological Tau accumulation in PS19 mice.

### CST treatment alters the neuronal populations and attenuates neuroinflammation in PS19 mice

To obtain an unbiased and high-resolution view of CST’s impact on cellular composition and transcriptional states in Tauopathy, we performed single-nucleus RNA sequencing (snRNA-seq) on hippocampal tissue from nTg, saline-treated PS19 (Sal/PS19), and CST-treated PS19 (CST/PS19) mice at 9 months of age the same cohort used for structural and biochemical analyses. After quality control, ∼35,000 nuclei were retained for downstream analysis (**Fig. 4A**). Unsupervised Leiden clustering of the snRNA-seq dataset identified 15 transcriptionally distinct clusters representing the major cellular constituents of the hippocampus along with their specific gene signatures (**S-Fig. 4A&B**). These included excitatory projection neuron populations such as CA1 Prosubiculum (CA1-ProS) and Subiculum-Prosubiculum (SUB-ProS) neurons, which contribute to hippocampal output and long-range cortical and subcortical connectivity, as well as CA3 and DG, CA2-IG-FC neurons involved in intrinsic hippocampal circuitry and information processing. Multiple inhibitory interneuron classes were also resolved, including Lamp5, Sncg, and Pvalb neurons, which regulate synaptic integration, oscillatory activity, and the balance of excitation and inhibition. Glial populations comprised astrocytes, which support synaptic function and metabolic homeostasis; microglia with perivascular macrophages (Micro-PVM), which mediate immune surveillance and neuroinflammatory responses; and oligodendrocyte precursor cells (OPCs), which contribute to myelin remodeling and axonal support. Additional smaller clusters represented other glial and vascular-associated cell types or transcriptionally less well-defined populations (**S-Fig.4A**).

**Figure 4.**
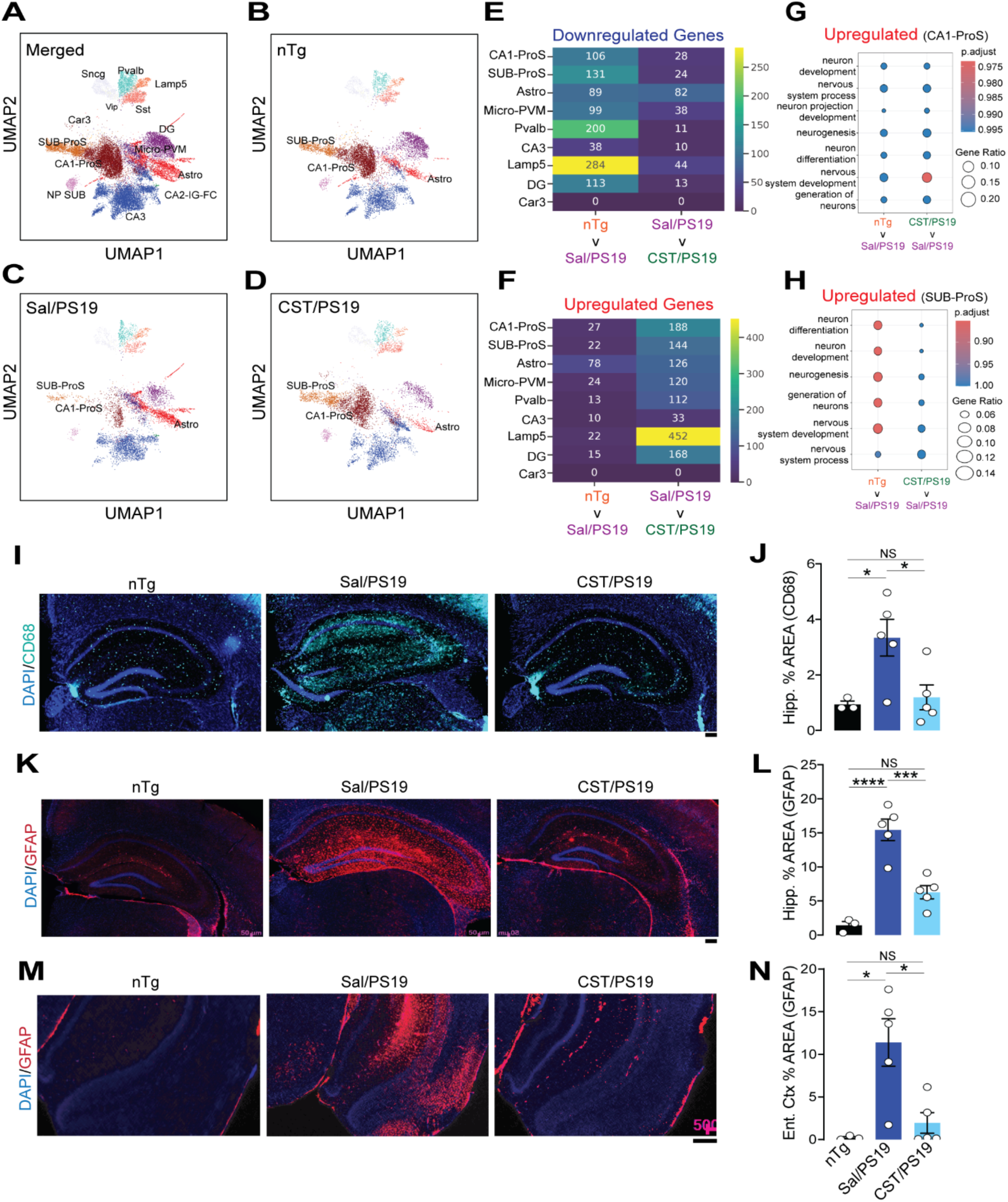
CST restores vulnerable neuronal populations and suppresses neuroinflammation in PS19 mice. (A–D) UMAP visualization of snRNA-seq data. (**A**) Merged dataset showing annotated neuronal and glial clusters. (**B**) Non-transgenic (nTg) brains display normal cellular distributions. (**C**) Sal/PS19 mice show loss of CA1-ProS and SUB-ProS neurons and expansion of Astro and Micro-PVM clusters. (**D**) CST treatment partially restores CA1-ProS/SUB-ProS populations and reduces Astro and Micro-PVM clustering. **(E&F)** Number of Downregulated and Upregulated differentially expressed genes (DEG) in the reference group (top) vs comparison group (bottom) for the identified neuronal and glial clusters. (**G**) Enriched upregulated pathways with GO terms in CA1-ProS cluster of nTg and CST/PS19 compared to Sal/PS19. (**H**) Enriched upregulated pathways with GO terms in SUB-ProS cluster of nTg and CST/PS19 compared to Sal/PS19. **(I&J)** CD68 immunostaining depicts increased hippocampal microglial activation in Sal/PS19 mice, which is significantly reduced by CST (nTg: n=3, Sal/PS19: n=5, CST/PS19: n=5). Scale bars = 200µM. **(K&L)** GFAP staining reveals robust hippocampal astrogliosis in Sal/PS19 mice; CST markedly decreases GFAP⁺ area (nTg: n=3, Sal/PS19: n=5, CST/PS19: n=5). Scale bars = 200µM. **(M&N)** GFAP staining in the entorhinal cortex also shows elevated astrocytosis in Sal/PS19 mice, partially rescued by CST (nTg: n=3, Sal/PS19: n=5, CST/PS19: n=5). Scale bars = 200µM. Data are mean ± SEM; Significance indicated as *p < 0.05, **p < 0.01, *\*\*\**p < 0.001, ****p < 0.0001; NS, not significant.

Cell-type proportion analysis showed significant remodeling of hippocampal cellular composition in PS19 mice, with partial restoration following CST treatment. Compared with Sal/nTg controls, Sal/PS19 samples exhibited a reduction in CA1-ProS and Sub-ProS excitatory neuronal clusters, accompanied by an expansion of astrocyte and Micro-PVM populations. CST treatment reversed these compositional shifts (**Fig. 4B-D, S-Fig.4C**), increasing the relative abundance of CA1-ProS and Sub-ProS neurons, thereby corroborating the hippocampal volumetric analysis (**Fig. 2B&C**) and reducing astrocyte and Micro-PVM clusters compared with Sal/PS19 mice. Additionally, we examined transcriptomic changes across the clusters. We found that, among neuronal clusters (CA1-ProS, SUB-ProS), expression of several genes is rescued by CST treatment, further supporting the beneficial effect of CST in these mice (**Fig. 4E&F**). Furthermore, pathway analysis of those genes in CA1-ProS and Sub-ProS showed that CST treatment upregulated several neuron-related biological processes, including neurogenesis, neuron differentiation, and neuron projection development, in PS19, comparable to nTg mice (**Fig. 4G&H**).

We further confirmed the glial transcriptomic signature through IHC. CD68, a marker of microglial phagocytic activation, was significantly higher in the hippocampus of saline-treated PS19 mice but was markedly lower in CST-treated PS19 mice and comparable to nTg levels (**Fig. 4I&J**). Similarly, glial fibrillary acidic protein (GFAP) immunoreactivity was drastically increased in both the hippocampus and entorhinal cortex of saline-treated PS19 animals but was markedly reduced in CST-treated mice (**Fig. 4K-N**). Since hippocampal and entorhinal neuroinflammation contribute to cognitive decline in Tauopathy, this reduction highlights CST’s neuroprotective effect.

We next quantified inflammatory cytokines in cortical lysates (**S-Fig. 5A-F**) and plasma (**S-Fig. 5G-L**). Saline-treated PS19 mice displayed robust elevations in IL-6, TNF-α, CXCL1, CXCL10, CXCL2, and CCL2 compared to nTg animals. CST treatment significantly reduced all four cytokines. These changes align closely with the IHC and snRNA-seq data, confirming suppression of neuroinflammation at the molecular level.

**Figure 5.**
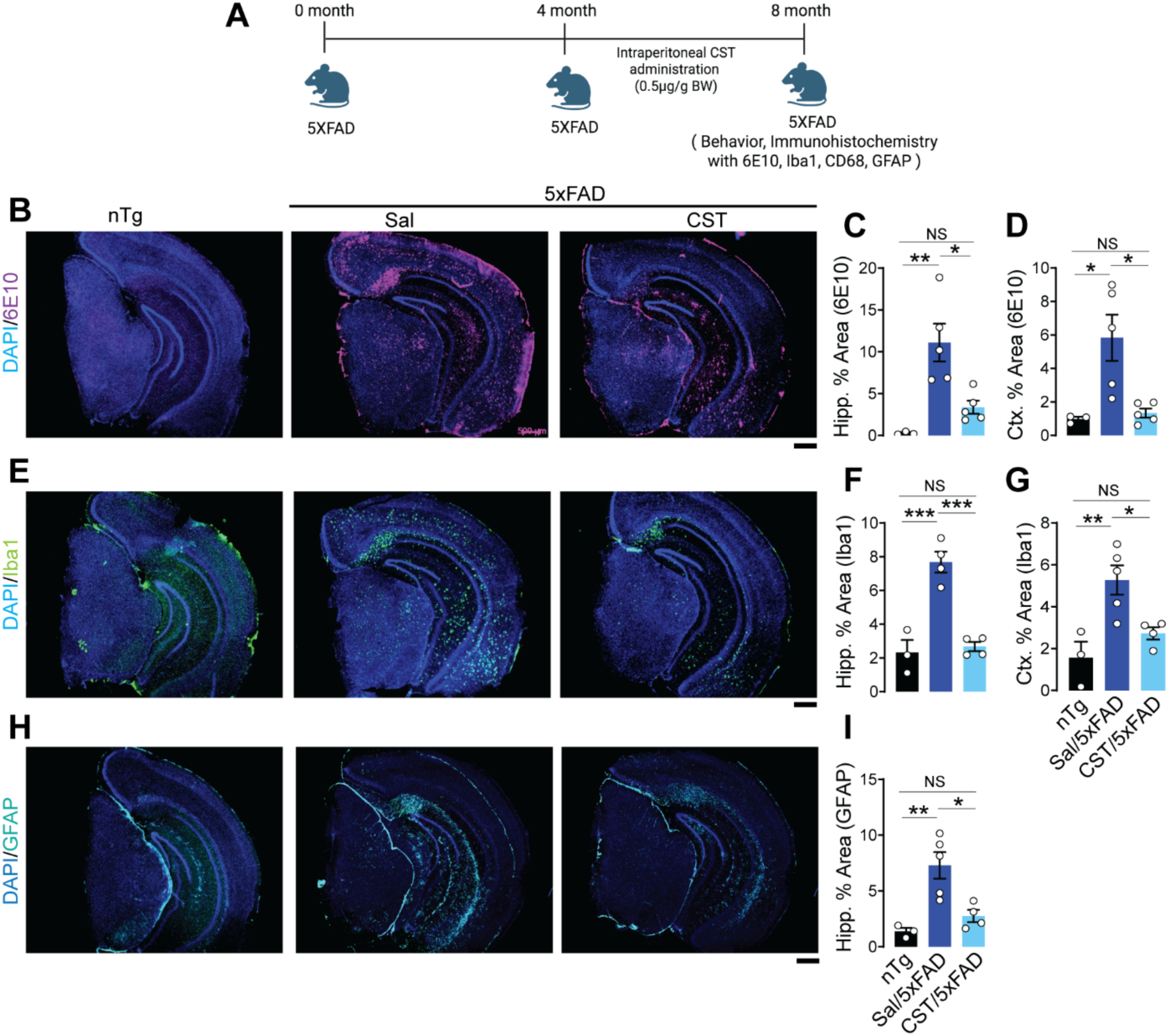
CST reduces amyloid plaque burden and neuroinflammation in 5xFAD mice. **(A)** Experimental design showing intraperitoneal CST administration (0.5 µg/g BW) to 5xFAD mice from 4 to 8 months of age, followed by behavioral and immunohistochemical analyses. **(B–D)** CST lowers amyloid plaque burden. 6E10 immunostaining **(B)** reveals extensive hippocampal and cortical amyloid deposition in saline-treated 5xFAD mice, whereas CST-treated mice exhibit significantly reduced 6E10⁺ area. Quantification of (**C**) hippocampal and (**D**) cortical regions confirms CST-mediated attenuation of amyloid load (nTg: n=3, Sal/5xFAD: n=5, CST/5xFAD: n=5). **(E–G)** CST decreases microgliosis. (**E**) Iba1 immunostaining shows pronounced microglial activation in saline-treated 5xFAD mice, with CST treatment markedly reducing Iba1⁺ area in both (**F**) hippocampus and (**G**) cortex (nTg: n=3, Sal/5xFAD: n=5, CST/5xFAD: n=5). **(H&I)** CST suppresses astrogliosis. (**H**) GFAP immunostaining demonstrates elevated astrocyte activation in saline-treated 5xFAD mice; (**I**) CST significantly reduces GFAP⁺ area in the hippocampus (nTg: n=3, Sal/5xFAD: n=5, CST/5xFAD: n=5). Data represent mean ± SEM; statistical significance determined by one-way ANOVA; significance indicated as *p < 0.05, **p < 0.01, *\*\*\**p < 0.001, ****p < 0.0001; NS, not significant. Scale bar = 500µM.

Taken together, these findings demonstrate that CST reshapes the composition of hippocampal cell types, restores neuronal populations, suppresses glial reactivity, and attenuates inflammatory cytokine level in PS19 mice. By integrating transcriptional, histological, and biochemical evidence, the data reveal a broad anti-inflammatory and neuroprotective effect of CST in Tauopathy.

### CST treatment reduces amyloid plaque burden and neuroinflammation in 5xFAD mice

Because AD is characterized by both Tau and amyloid pathology ^39,40^, we next examined whether CST could modulate Aβ plaque formation and associated neuroinflammation in an amyloid-driven model. We used the 5xFAD mouse line (Tg(APPSwFlLon, PSEN1*M146L*L286V)6799Vas/Mmjax), which develops rapid and robust Aβ deposition. CST treatment (0.5 µg/g body weight, i.p., 3x/week) was initiated at 4 months of age, before overt plaque formation, and continued until 8 months, at which point brains were harvested for neuropathological analysis (**Fig. 5A**).

Amyloid plaque burden was quantified using 6E10, an antibody recognizing the Aβ1–16 epitope within the amyloidogenic core. Saline-treated 5xFAD mice exhibited extensive 6E10-positive plaque deposition in both hippocampus and cortex. In contrast, CST-treated 5xFAD mice showed a marked reduction in 6E10 staining across both regions (**Fig. 5B–D**), indicating that CST attenuates amyloid accumulation *in vivo*.

Given that 5xFAD pathology is accompanied by early and aggressive neuroinflammation ^41,42^, we next assessed microglial and astrocytic activation. Immunohistochemical analysis for Iba1 revealed pronounced microgliosis in saline-treated 5xFAD mice, whereas CST significantly reduced the Iba1-positive area in both the hippocampus and the cortex (**Fig. 5E–G**). Similarly, CD68 intensity was diminished in CST-treated animals (**S-Fig. 6A-C**). Astrocytic reactivity was evaluated using GFAP. Saline-treated 5xFAD mice displayed extensive GFAP immunoreactivity in the hippocampus and entorhinal cortex, whereas CST treatment resulted in a robust decrease in GFAP-positive area (**Fig. 5H&I**).

**Figure 6.**
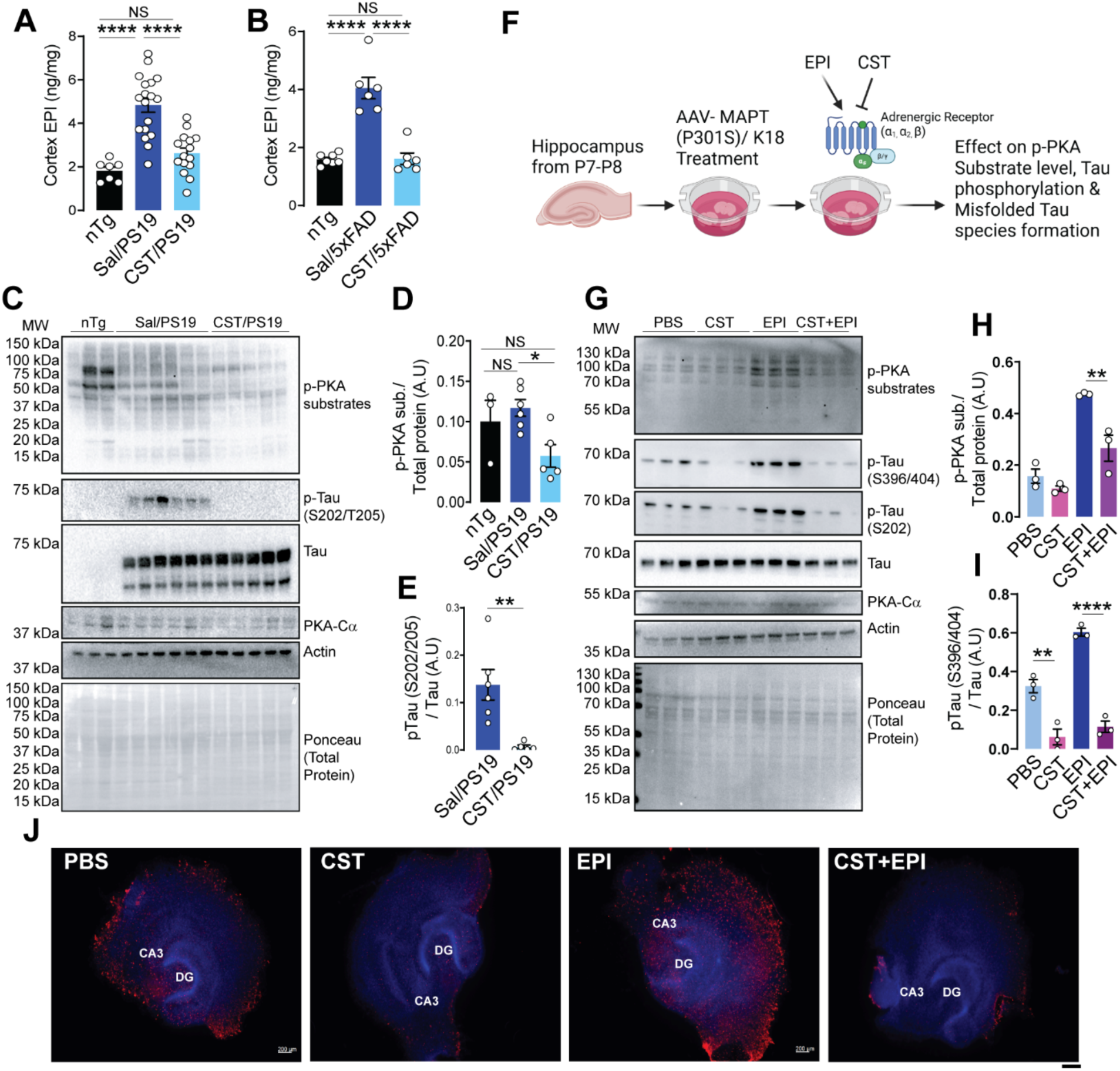
CST inhibits adrenergic/PKA hyperactivation and Tau phosphorylation in PS19 mice and an AAV-MAPT (P301S/K18) model. **(A)** Cortical epinephrine (EPI) levels are markedly elevated in Sal/PS19 compared with nTg controls, and CST treatment significantly lowers EPI concentrations (nTg: n=7, Sal/PS19: n=18, CST/PS19: n=15). **(B)** Cortical EPI levels are markedly elevated in Sal/5xFAD compared with nTg controls, and CST treatment significantly lowers EPI concentrations (nTg: n=7, Sal/5xFAD: n=6, CST/5xFAD: n=6). **(C–E)** CST suppresses pathological PKA signaling in the PS19 hippocampus. (**C**) Immunoblots show increased phospho-PKA substrates and elevated p-Tau (S202/T205) in Sal/PS19 mice; CST treatment reduces both, while total Tau and PKA-Cα remain unchanged. (**D&E**) Quantification confirms CST-mediated reduction of PKA activity and Tau phosphorylation (nTg: n=3, Sal/PS19: n=6, CST/PS19: n=5). **(F)** Schematic illustrating CST blockade ADR–driven PKA activation. In AAV-MAPT (P301S/K18)–transduced hippocampal slices, EPI strongly activates PKA signaling and enhances Tau phosphorylation, whereas CST antagonizes adrenergic/PKA hyperactivation and prevents formation of misfolded Tau species. **(G–I)** CST counteracts EPI-induced PKA activation and Tau pathology. (**G**) Immunoblot analysis shows that EPI markedly increases phospho-PKA substrates and p-Tau species (S202, S396/404), CST alone has minimal effect on PKA activity. Still, CST prevents EPI-induced PKA hyperactivation (PBS: n=3, CST: n=3, EPI: n=3, CST+EPI: n=3). (**H&I**) Quantification demonstrates CST’s ability to blunt adrenergic/PKA overactivation and its downstream Tau phosphorylation signatures. **(J)** Immunofluorescence staining (CA3, DG) reveals substantial accumulation of misfolded Tau in EPI-treated slices, which is significantly reduced by CST and nearly abolished by CST+EPI (PBS: n=3, CST: n=3, EPI: n=3, CST+EPI: n=3). Scale bar = 200µM. A.U.: Arbitrary Unit. Data shown as mean ± SEM; significance indicated as *p < 0.05, **p < 0.01, *\*\*\**p < 0.001, ****p < 0.0001; NS, not significant.

Collectively, these results demonstrate that CST not only reduces Aβ plaque burden but also suppresses glial activation in the 5xFAD model.

### CST treatment reduces EPI-mediated PKA activation Tau phosphorylation

We previously reported that elevated EPI levels and alpha-adrenergic receptor (ADRA) activation can drive Tau phosphorylation and aggregation ^8^. In contrast, norepinephrine (NE) levels do not show substantial changes across disease stages ^8^. From transcriptomic analysis, we observed an imbalance in ADR between Sal- and CST-treated PS19 (**S-Fig. 7A**) and we further confirmed that by qPCR (**S-Fig. 7B**). Given that CST is a potent inhibitor of chromaffin cell catecholamine release ^11,12,43,44^, including nicotine-evoked EPI secretion from primary hippocampal neurons ^45^, we next evaluated whether chronic CST treatment lowers brain EPI levels in Tauopathy and amyloid models.

To assess endogenous catecholamine levels, we quantified EPI and NE in cortical extracts from nTg, saline-treated PS19, CST-treated PS19, saline-treated 5xFAD, and CST-treated 5xFAD mice using Ultra Performance Liquid Chromatography (UPLC). In both PS19 and 5xFAD mice, EPI levels were elevated and is reduced by CST treatment back to nTg levels (**Fig. 6A&B**). NE levels also showed a similar trend (**S-Fig. 7C&D**).

These results suggest that CST restrains pathological elevations in catecholamines. Interestingly, transcriptomic analysis revealed increased alpha-1 ADRA (ADRA1) levels and decreased alpha-2 ADRA (ADRA2) levels in Sal/PS19 compared with CST/PS19.

Because heightened EPI signaling activates PKA - a serine/threonine kinase known to phosphorylate multiple cellular protein ligands and affects the Tau phosphorylation downstream at multiple pathological epitopes - we next assessed PKA activity in CST- and saline-treated PS19 mice. Immunoblotting using a phospho-PKA substrate (RRXS/T) antibody revealed an increase in PKA-dependent substrate phosphorylation in saline-treated PS19 mice cortex, which is markedly reduced after CST treatment (**Fig. 6C&D**). The abundance of the PKA catalytic subunit remains unaltered, indicating that CST normalizes ADR/PKA activity and downstream signaling cascade rather than reducing PKA expression. In parallel, CST also lowered pTau (S202/T205) levels in PS19 cortex as we observed earlier (**Fig. 6E**), reinforcing the link between EPI-PKA activation and Tau hyperphosphorylation.

To investigate whether CST blocks EPI-driven Tau pathology, we performed hippocampal OTSC experiments with four conditions: PBS, CST, EPI, and CST + EPI (**Fig. 6F**). Consistent with previous reports, EPI treatment markedly increased PKA substrate phosphorylation, Tau phosphorylation, and MC1-positive misfolded Tau accumulation compared to PBS controls (**Fig. 6G-J**). CST alone reduced both pTau and MC1 staining, confirming its protective effect in an *ex vivo* system. Importantly, co-treatment with CST and EPI significantly attenuated the EPI-induced increase in Tau phosphorylation, aggregation, and phospho-PKA substrate levels **(Fig. 6G-J, S-Fig. 7E**). Meanwhile, total PKA catalytic subunit abundance remained unchanged across all groups. These data demonstrate that CST interferes with EPI-mediated PKA activation and its downstream effects on Tau pathology.

Together, these results mechanistically link CST’s neuroprotective activity to its ability to suppress EPI-driven adrenergic stress. By reducing EPI levels, blunting PKA activation, and preventing Tau phosphorylation and misfolding, CST interrupts a pathogenic cascade that contributes to neurodegeneration in both Tauopathy and amyloid-driven models.

## Discussion

Tauopathies, including AD, CBD, and PSP, arise from convergent pathogenic processes encompassing Tau hyperphosphorylation ^46–48^, impaired proteostasis ^49,50^, synaptic dysfunction ^51,52^, and chronic neuroinflammation ^4,5^. In AD, extracellular Aβ plaques and intracellular neurofibrillary tangles synergistically promote neuronal loss and cognitive decline ^39,40^. While kinase dysregulation ^53^, adrenergic stress ^54^, metabolic dysfunction ^55^, and glial activation ^56^ are recognized contributors to disease progression, neuroendocrine regulation of these pathways has remained poorly defined.

CgA, stored in DCVs, is elevated in AD CSF and associates with proteinopathic aggregates in postmortem brain ^8–10^. We demonstrate that Tauopathies are characterized by a marked reduction in the neuroprotective CgA-derived peptide CST, accompanied by a reciprocal increase in PST, in human AD, CBD, and PSP. Similar alterations have been reported in metabolic disorders including type 1 diabetes ^37^, type 2 diabetes (T2D) ^57^, and hypertension ^17,38^. CST deficiency was recapitulated in PS19 mice, indicating that loss of CST is a conserved feature across Tauopathies. In contrast, PST - a functional antagonist of CST ^37^ - was elevated in both AD and T2D tissues ^57^, revealing a previously unrecognized neuroendocrine imbalance linking metabolic stress to neurodegeneration. Mechanistically, CST supplementation exerted robust neuroprotective effects across *in vitro*, *ex vivo*, and *in vivo* models. CST reduced Tau phosphorylation at multiple pathological epitopes, suppressed misfolded Tau accumulation (MC1), and attenuated Tau seeding activity - key drivers of Tau propagation and toxicity ^58,59^. In PS19 mice, CST preserved hippocampal structure and improved cognitive, working memory, and motor performance, consistent with CST’s established neuroprotective roles in hypertensive rat model ^60^.

At the systems level, single-nucleus RNA-seq revealed that Tauopathy induces coordinated remodeling of the hippocampal cellular landscape, marked by loss of CA1-ProS and Sub-ProS neurons and expansion of reactive astrocytes and activated microglia/perivascular macrophages (Micro-PVM). This pattern reflects a feed-forward neuron–glia inflammatory loop that accelerates neurodegeneration ^56,61^. CST treatment normalized this pathological remodeling, restoring vulnerable neuronal populations while suppressing reactive glial states. Concordantly, CST reduced expression of CD68, GFAP, IL-6, TNF-α, CXCL1, and CXCL10, extending its known peripheral anti-inflammatory actions ^20,38^ to the central nervous system and establishing CST as a regulator of neuroimmune homeostasis in Tauopathy.

CST also reduced Aβ plaque burden and gliosis in 5xFAD mice, supporting a broader role in stress-responsive neurodegenerative pathways. Given that Aβ pathology promotes adrenergic hyperactivation ^8,62^, metabolic stress, and glial priming ^56^, these findings indicate that CST modulates shared upstream mechanisms linking amyloid and Tau pathologies.

A key mechanistic insight from this study is that CST suppresses EPI levels and restrains EPI-driven PKA signaling, a pathway strongly implicated in stress-induced Tau phosphorylation, synaptic dysfunction, and neuronal vulnerability ^63–65^. ADR activation elevates cAMP and PKA activity, promoting Tau phosphorylation at epitopes that destabilize microtubules and facilitate misfolding and aggregation ^66,67^. Elevated PKA activity in human AD brain correlates with Tau pathology and cognitive impairment. CST inhibits EPI release from both adrenal medulla ^43^ and hippocampal neurons ^45^, directly linking neuropeptide imbalance to pathological ADR signaling. Importantly, CST reduced PKA substrate phosphorylation without altering PKA catalytic subunit levels, indicating normalization of stress signaling rather than nonspecific kinase inhibition. CST also blocked EPI-induced Tau phosphorylation and misfolding in hippocampal slice cultures, providing direct mechanistic evidence that adrenergic restraint protects against Tauopathy. Considering hypertension as a major risk factor for neurodegeneration ^68^, these findings are highly significant.

Collectively, these findings establish CST as an endogenous neuroendocrine regulator that integrates adrenergic, inflammatory, and metabolic pathways upstream of Tau and amyloid pathology. Unlike therapeutic strategies targeting single disease components—such as anti-Aβ antibodies ^27^, anti-Tau vaccines ^29^, or kinase inhibitors ^22–24,26,69^ - CST engages convergent mechanisms that drive neurodegeneration. This multimodal action highlights CST and its analogs as promising translational candidates for AD, Tauopathies, and related proteinopathies.

## Materials and Methods

### Animals

All animal studies were approved by the Institutional Animal Care and Use Committees (IACUC) of the University of California, San Diego (UCSD) and the Veterans Affairs San Diego Healthcare System, and were conducted in accordance with relevant National Institutes of Health (NIH) guidelines. Transgenic PS19 mice (Tau P301S; B6;C3-Tg(Prnp-MAPT*P301S)PS19Vle/J)* and *5xFAD mice (B6.Cg-Tg(APPSwFlLon,PSEN1*M146L*L286V)6799Vas/Mmjax) were purchased from The Jackson Laboratory (Bar Harbor, ME, USA). Heterozygous PS19 mice were bred in house and used for all experiments, whereas 5xFAD mice were used directly following purchase from The Jackson Laboratory.

Both male and female mice were included in all experimental groups. Sex was considered a biological variable in the experimental design and data analysis, and no animals were excluded based on sex. Lyophilized CST peptide (SSMKLSFRARAYGFRGPGPQL) from GenScript was dissolved in saline and administered intraperitoneally to the animals. Animals were housed under controlled conditions with a 12-hour light/12-hour dark cycle and had ad libitum access to food and water. Mice were maintained on a standard normal chow diet (NCD; 14% kcal from fat; LabDiet 5P00).

### Genotyping

Mice were ear-tagged for identification, and tail snips were collected for genotyping. Genomic DNA was isolated from tail tissue using the AccuStart™ Genotyping Kit and amplified by PCR using AccuStart™ GelTrack™ PCR SuperMix according to the manufacturer’s instructions. PS19 genotyping was performed using the following primers: forward primer (common to wild-type and mutant alleles), 5′-TTG AAG TTG GGT TAT CAA TTT GG-3′; reverse primer (wild-type), 5′-TTC TTG GAA CAC AAA CCA TTT C-3′; and reverse primer (mutant), 5′-AAA TTC CTC AGC AAC TGT GGT-3′. 5xFAD genotyping was performed with the following primers: Common Forward: 5’-ACC CCC ATG TCA GAG TTC CT-3’, Wild-type Reverse: 5’-TAT ACA ACC TTG GGG GAT GG-3’, Mutant Reverse: 5’-CGG GCC TCT TCG CTA TTA C-3’.

### Tissue extract preparation and Western Blotting

Tissue extraction and western blotting was done according to previously published work ^8^. 15mg tissue was incubated in RIPA buffer (150mM NaCl, 1% Triton-X, 12mM Na-deoxycholate, 0.1% SDS, 50mM Tris pH 8.0, 25mM EDTA, 1mM PMSF, 5% Glycerol, 50mM NaF, 1mM Na3VO4, 1mM Na4P2O7, 25mM β-Glycerophosphate, 50mM DTT and PIC) for 10min. A handheld motor homogenizer was used to homogenize the tissue, followed by centrifugation at 12,500 rpm for 30 min at 4°C. The supernatant was collected, and protein concentration was determined using Bradford Protein Assay Reagent. 15μg protein was loaded in each lane of a 12% Tris-Glycine SDS Gel, and post-running protein was transferred to a PVDF membrane, followed by blocking with 5% BSA for 2 hours. Primary antibody incubation was done at 4°C.

### Hippocampal Organotypic Slice Culture (OTSC)

Organotypic brain slice cultures (BSC) were prepared using hippocampal slices from wild-type P8-10 pups, following previously established methods with some modifications. Pups were euthanized by decapitation, and the bilateral hippocampi were quickly dissected in oxygenated aCSF (125mM NaCl, 2.4mM KCl, 1.2mM NaH2PO4, 1mM CaCl2, 2mM MgCl2, 25mM NaHCO3, and 25mM Glucose). The aCSF had been oxygenated beforehand. The hippocampi were sliced into 300 μm sections with a tissue chopper and cultured in Millicell inserts (Millipore PICM0RG50) placed in 6-well plates, with five slices per insert. The culture medium was BSC media comprising 50% (v/v) Basal Medium Eagle (BME), 25% (v/v) Heat-Inactivated Horse Serum, 1% (v/v) Glutamax, 0.5% (v/v) Pen/Strep, 0.033% (v/v) insulin, 45mM D-Glucose, and 25mM HEPES in EBSS buffer, all sterile-filtered through a 0.2μm filter.

On DIV0, slices were transfected with AAV2-hTauP301S (ViroTek) at a titer of 2E10 mg/mL. On DIV2, the slices were treated with 1.5 μg/mL of homemade K18-PFF ^8^. On DIV4, 2µM CST or PST was added to the media, which was refreshed during each media change every 2-3 days. On DIV14, slices were either harvested for western blot by extraction in RIPA or fixed in 4% PFA at room temperature for 2-3 hours. After fixation, slices were permeabilized overnight in 1% Triton X-100 in PBS. Antigen retrieval was performed using citrate buffer (pH 6.0) in a pressure cooker for 15 minutes at low pressure. Slices were then blocked in 20% BSA with 0.4% Triton X-100 in PBS for 2-3 hours at room temperature. The primary antibody was incubated in 0.4% Triton X-100 in PBS for 48 hours at 4°C. Subsequently, slices were washed six times for 10 minutes each in PBS, then incubated with fluorophore-conjugated secondary antibodies and DAPI for 2 hours at room temperature. Final washing involved six additional 10-minute PBS washes, after which slices were mounted on glass slides with their membrane inserts and cover slipped using Fluoromount-G. Images were obtained using a Keyence BZX-700 fluorescent microscope.

### Immunohistochemistry

Mice were anesthetized with isoflurane, then transcardially perfused. Perfusion was performed with phosphate-buffered saline (PBS). Brains were then dissected and placed in zinc-formalin (Zn-Formalin). Brains were fixed in Zn-Formalin for 48 hours. Subsequently, Zn-Formalin was replaced with a 30% sucrose solution. Samples were kept in 30% sucrose for 72 hours at 4°C. Coronal sectioning was then performed. Sections of 30 μm thickness were prepared using a sliding freezing microtome (Epredia). Sections were stored at −20°C in cryoprotectant solution. For each animal, 7 to 8 sections were collected, spanning the anterior to posterior regions of the hippocampus. Sections were thoroughly washed to remove excess cryoprotectant. A series of six 10-minute washes with 1X PBS was performed, followed by incubation with the primary antibody. Incubation was carried out in PBS containing 0.4% Triton-X for 24 hours at 4°C. After incubation, the primary antibody was removed. Sections were washed three times for 15 minutes each with 1X PBS. Subsequently, secondary antibody and DAPI (1:2000) were added. The sections were incubated at room temperature for 1 hour, followed by three 15-minute washes with PBS. Next, sections were mounted on glass slides using Fluoromount G. Stained slides were imaged using a Keyence fluorescence microscope.

### CST and PST Enzyme Immunoassays

CST and PST levels were quantified in mouse plasma, mouse cerebral cortex, and human cortical tissue using commercially available species-specific enzyme immunoassay (EIA/ELISA) kits. Mouse CST concentrations were measured using a mouse CST EIA kit (RayBiotech Life, Peachtree Corners, GA, USA), and mouse PST levels were determined using a mouse PST EIA kit (MyBioSource, Inc., San Diego, CA, USA). Human CST and PST levels were quantified using corresponding human-specific ELISA kits in accordance with the manufacturers’ protocols.

For tissue measurements, cortical samples were homogenized in appropriate assay buffer, clarified by centrifugation, and the resulting supernatants were used for analysis. CST levels in cortical tissue were expressed as nanograms per milligram of tissue (ng/mg tissue). Plasma samples were assayed directly or following dilution as required to remain within the linear range of the standard curves, and plasma CST concentrations were expressed as nanomolar (nM). All samples and standards were assayed in duplicate, absorbance was measured using a microplate reader at the recommended wavelength, and peptide concentrations were calculated from standard curves generated for each assay. Samples were processed in parallel to minimize inter-assay variability.

### Single-nucleus RNA-sequencing (snRNA-seq)

*Nucleus Isolation, Library Preparation and Sequencing*: Hippocampal regions from nTg, Sal/PS19, and CST/PS19 were used, with hippocampus from four different animals in each group pooled together for each experimental set. Nuclear isolation was performed using three buffers: Nuclear Isolation Medium 1 (NIM 1: 250 mM sucrose, 25 mM KCl, 5 mM MgCl2, 10 mM Tris pH 8. 8.0), Nuclear Isolation Medium 2 (NIM 2: 0. 95 parts NIM 1, 1 µM DTT, PIC), and Homogenization Buffer (HB: 0. 97 parts NIM 2, RNaseIN 0. 4 U/µl, SuperaseIn 0. 2 U/µl, 0.1% Triton X- 100), prepared according to a previously standardized protocol ^70^. A prechilled Dounce homogenizer was filled with an appropriate amount of HB. The hippocampal sections, cut into 2-3 mm³ pieces in prechilled homogenization buffer, were transferred to a Dounce homogenizer, where the tissue was homogenized 7 times and kept on ice for 3 minutes. The homogenate was strained through a 70 µm strainer (BD Falcon). A 10µl aliquot was taken and loaded onto a hemocytometer with Trypan Blue to assess nuclear integrity and isolation. To the remaining strained sample, 10 mM Ribonucleoside Vanadyl Complex (New England Biolabs) was added, and the mixture was incubated for 1 minute. The samples were then centrifuged at 200g for five minutes in a swing-bucket rotor. The supernatant was removed, and 500 µl of solution containing DPBS, 10 mM VRC, DNase I, and RNase inhibitor was added to the pellet. The pellet was gently pipetted multiple times to mix and strained through a 35 µm strainer (BD Falcon). The mixture was centrifuged again at 200g for 5 minutes, the supernatant was removed, and the pellet was dissolved in 200µl of DPBS. The solution was mixed thoroughly. Nuclei were stained with Hoechst dye and checked for integrity and debris. One million nuclei per sample were taken and fixed using Parse BioSciences Evercode ™ Nuclei Fixation V 3 for further analysis (Parse Biosciences, Seattle, WA). Fixed nuclei were stored at -80°C. On the day of the experiment, they were thawed on ice, and the integrity of the nuclei and count were checked under the microscope. The samples were processed according to the manufacturer’s protocol for the Parse BioSciences Evercode ™ NTG v 3 kit. The kit uses a combination of barcodes, and all samples were prepared together with five other samples from a different experiment and divided into eight sublibraries. These sublibraries were quantified and assessed for quality using Qubit and then sequenced.

*Single-cell RNA-seq data preprocessing:* Single-cell RNA-seq data were first analyzed using Trailmaker from Parse Biosciences. The downloaded data from Trailmaker is analyzed using Scanpy (v1.9.3). Cells with a doublet score ≥ 0.1 were removed to exclude potential doublets. Further quality control filters removed cells with fewer than 4,000 detected features (nFeature_RNA < 4000) or with high mitochondrial gene content (percent.mt < 4). Total-count normalization was performed to a target sum of 10,000 transcripts per cell using sc.pp.normalize_total, followed by log1p transformation (sc.pp.log1p). Highly variable genes (HVGs) were identified using sc.pp.highly_variable_genes with parameters min_mean = 0.0125, max_mean = 3, and min_disp = 0.5, and visualized using sc.pl.highly_variable_genes. Only HVGs were retained for downstream UMAP clustering analysis, and the raw unfiltered expression matrix was stored in raw data. Technical covariates, including total RNA counts (nCount_RNA) and mitochondrial gene percentage (percent.mt), were regressed out using sc.pp.regress_out, followed by z-score scaling with a maximum value of 10 (sc.pp.scale).

*Dimensionality reduction, Clustering, and cell-type annotation:* Principal component analysis (PCA) was performed using sc.tl.pca with the ‘arpack’ solver. PCA embeddings were visualized, and variance explained was assessed using sc.pl.pca and sc.pl.pca_variance_ratio. Neighborhood graphs were computed with n_neighbors = 10 and n_pcs = 40 (sc.pp.neighbors), and two-dimensional embeddings were generated using UMAP (sc.tl.umap) and t-SNE (sc.tl.tsne). Graph-based clustering was performed using the Leiden algorithm (sc.tl.leiden) with resolution = 0.9, random_state = 0, n_iterations = 2, and undirected graphs. Cluster assignments were visualized on UMAP embeddings along with marker gene expression. For automated cell-type annotation, the CellTypist framework (v1.6.3) was used. The model Mouse_Isocortex_Hippocampus ^71^ was loaded via celltypist.models.Model.load, and annotations were assigned to cells using celltypist.annotate with majority voting. Predicted labels were integrated into the AnnData object for downstream stratified analyses.

*scVI model setup and training:* Single-cell transcriptomes were modeled using scVI-tools (v0.20.3), a deep generative framework based on variational autoencoders ^72^. The AnnData object was configured using SCVI.setup_anndata, with sample identity included as a categorical covariate, and library size, number of detected genes, mitochondrial gene percentage, and doublet score included as continuous covariates to account for technical variation ^73^. A scVI model was instantiated with default architecture and optimization parameters and trained on the filtered dataset. Automatic hardware selection enabled acceleration on available computational resources. After training, the model was saved and used exclusively for downstream inference and differential expression (DE) analyses without additional fine-tuning. Cells were assigned to major cell types using a majority-voting annotation using celltypist.annotate as described before. These cell-type annotations stratified downstream analyses. Experimental conditions were defined by group labels (A, B, and C), and only cell types present in both comparison groups were included in DE analyses.

*Cell-type–specific differential expression analysis:* DE analysis was performed using scVI’s Bayesian differential expression framework ^72^. Pairwise comparisons were conducted for each cell type between experimental groups (A vs B, B vs C, and A vs C).

Boolean indexing was used to select cells from the groups of interest; comparisons were skipped for cell types lacking representation in either group. DE was estimated using scVI’s differential_expression function, which computes posterior distributions of gene-wise log fold changes while accounting for technical noise and latent biological variability. Results for all cell types were concatenated and stored as separate tables for downstream analysis. Genes were considered significantly differentially expressed if they satisfied all of the following criteria: (1) scVI-derived false discovery rate threshold (is_de_fdr_0.05), (2) absolute median log fold change > 1, and (3) detectable expression in ≥1% of cells in either group. Upregulated (log fold change < -1) and downregulated (log fold change > 1) genes were analyzed separately.

*Aggregation of differential expression across cell types:* For each comparison, the number of significant DE genes per cell type was quantified. A unified set of observed cell types was used to construct DEG count matrices, with missing values filled as zeros. DEG count tables from all pairwise comparisons were merged into a single matrix indexed by cell type, enabling direct comparison of DE burden across conditions. Heatmaps of significant DE gene counts per cell type were generated using Seaborn (v0.12.2) with consistent color scales and annotated counts. Separate heatmaps were produced for upregulated and downregulated genes. Figures were rendered at high resolution (300–400 dpi) and formatted for publication-quality output.

*Software and computational environment:* Analyses were performed in Python using NumPy (v1.22.4) and pandas (v2.0.0) for computation and data handling, Scanpy (v1.9.3) for single-cell preprocessing, the CellTypist framework (v1.6.3) for cell annotation, scVI-tools (v0.20.3) for probabilistic modeling and DE analysis, and Seaborn (v0.12.2) and Matplotlib for visualization. All software versions were recorded to ensure reproducibility. Analyses were performed on a multi-GPU system with CUDA (v11.1) enabled with 4 GPU devices (GeForce GTX 1080 Ti). Codes are released on GitHub: sahoo00/snRNASeq-PS19.

*Pathway analysis:* We performed Gene Ontology (GO) enrichment analysis to identify biological processes associated with differentially expressed genes in single-nucleus RNA-sequencing data. For each cell population and condition, we created lists of We analyzed differentially expressed genes and examined upregulated genes separately. Gene identifiers were mapped to Entrez Gene IDs using the org.Mm.eg.db annotation package in R. GO enrichment analysis was performed with the clusterProfiler package (v4.16.0), focusing on the Biological Process category. A hypergeometric test assessed enrichment, and p-values were adjusted for multiple comparisons using the Benjamini–Hochberg false discovery rate (FDR) method. For comparisons across conditions, we utilized the compareCluster function for integrated enrichment analysis and visualized the results as dot plots using the Enrichplot package.

### Measurement of catecholamines

Catecholamine levels in cortical tissue was quantified by Ultra Performance Liquid Chromatography (UPLC) coupled with electrochemical detection (HPLC-ECD) as described previously ^8^. Samples were separated on an Atlantis dC18 column (100 Å, 3 μm, 3.0 × 100 mm; Waters Corp., Milford, MA) using an ACQUITY UPLC H-Class System (Waters Corp.) equipped with an electrochemical detector (ECD model 2465; Waters Corp.). The mobile phase was delivered isocratically at a flow rate of 0.3 mL/min and consisted of a 95:5 (v/v) mixture of phosphate–citrate buffer and acetonitrile. Tissue samples were homogenized in 0.1 N HCl, and 3,4-dihydroxybenzylamine (DHBA; 400 ng) was added as an internal standard. Catecholamines were adsorbed onto approximately 15 mg of activated aluminum oxide with gentle rotation for 10 min. The aluminum oxide was washed with 1 mL of distilled water, and catecholamines were eluted with 100 μL of 0.1 N HCl. Total protein concentration in cortical homogenates was determined using a bicinchoninic acid (BCA) protein assay, and catecholamine levels were normalized to protein content. The electrochemical detector was set at 200 pA and the chromatographic data were acquired and analyzed using Empower software (Waters Corp.). Catecholamine concentrations were calculated using the internal standard method and normalized to DHBA recovery. Cortical catecholamine levels were expressed as nanograms per milligram of protein (ng/mg protein).

### Measurement of Cytokines

Cortical tissues from nTg, Sal/PS19 and CST/PS19 mice were taken. Samples were homogenized in PBS and centrifuged at 12,500 RPM for 30 minutes at 4 °C. The resulting supernatant was collected for ELISA assays. Cytokine concentrations were measured using the U-PLEX mouse cytokine assay kit (Meso Scale Diagnostics, Rockville, MD) according to the manufacturer’s protocol, with detection performed on a MESO SECTOR S 600MM Ultra-Sensitive Plate Imager. Cytokine levels were expressed as pg per mg of protein. Plasma cytokines (25 μl) were measured using the U-PLEX mouse cytokine assay kit (Meso Scale Diagnostics, Rockville, MD).

### Behavior Studies

*Y-maze:* Mice were tested in the Y-Maze (ConductScience) following a previously published method ^74^, with some modifications. Each animal was allowed to explore the maze freely for 10 minutes. We used an automated system (ConductScience ) to record the number of arm entries and reentries, as well as the distance traveled. A spontaneous alternation was counted when a mouse entered a different arm on each of three consecutive entries. The percentage of spontaneous alternation was calculated as ((number of spontaneous alternations)/(net arm entries)) × 100.

*Two-Trial Memory Test:* The procedure was performed as previously described ^75^. The two-trial Y maze test incorporated visual cues positioned at the end of each arm. Sixteen hours after the training session, during which the novel arm remained closed, the mice were returned to the Y maze and allowed to explore all arms without restriction for 5 minutes. The time spent in both the novel and familiar arms was recorded using ConductScience software, and the ratio of time spent in the novel arm to that in the familiar arm was calculated.

*Grip Strength:* Forelimb (front two paws) and all-limb (four paws) grip strength was assessed using the Grip Strength Meter. Following the manufacturer’s protocol, each mouse was held by the tail, lowered toward the apparatus, and permitted the metal grid to grasp with either two or four paws. Mice were then pulled backward in a horizontal direction, and the force exerted on the grid immediately before grip loss was recorded as peak tension, which was converted to grams by the transducer. For each mouse, peak force was measured twice in succession for both the forelimbs and all limbs. The mean of the two trials was used for analysis. A minimum interval of five minutes was provided between trials for each mouse.

*Rotarod:* The rotarod test was conducted to assess motor function. In this test, a horizontal rod rotates around its axis, requiring mice to coordinate their movements to remain on the apparatus. The mice must coordinate with the rod’s movement to avoid falling. The duration that each mouse stays on the rod is recorded as a measure of motor performance. On day 1, each mouse undergoes five trials on the rotarod. On day 2, each mouse undergoes seven trials, and the time from each trial is recorded. The recorded times from each trial are plotted and compared to evaluate changes in motor function.

*Nesting:* Nest-building behavior was used to assess cognitive function in mice. PS19 and wild-type (NTG) mice were housed individually and provided with intact compressed cotton nestlets placed in the center of each cage. After 24 hours, images of each nest were captured. Nest quality was evaluated using a 1–5 scale, as previously described. The scoring criteria were as follows: 1, untouched nestlet; 2, partially torn nestlet; 3, partially complete nest; 4, nest almost formed; and 5, complete nest. All mice underwent testing after six weeks of treatment.

### Nissl Staining, Volumetric Analysis, and Thickness Measurement of DG and CA1

Nissl staining was conducted to visualize neuronal cell bodies and Nissl substance in fixed tissue sections. Brain or spinal cord tissues were fixed in 4% paraformaldehyde prepared in 0.1 M phosphate buffer or phosphate-buffered saline. The fixed tissues were sectioned using a cryostat or vibratome at 30μm thickness. Sections were mounted onto positively charged microscope slides and air-dried overnight on a slide warmer to ensure adequate adhesion.

A 0.1% cresyl violet staining solution was used for this purpose. Immediately before use, 10 drops (approximately 0.3 mL) of glacial acetic acid were added, and the solution was filtered. Mounted sections were defatted by immersion in a 1:1 mixture of ethanol and chloroform overnight, then rehydrated through 100% and 95% ethanol to distilled water. To prevent tissue detachment, frozen sections were not placed directly into water.

Sections were stained in 0.1% cresyl violet solution for 5–10 minutes. The staining solution was prewarmed to 37 °C to enhance dye penetration and achieve uniform staining. After staining, sections were briefly rinsed in distilled water and differentiated in 95% ethanol for 2–30 minutes, with microscopic monitoring to ensure optimal contrast. Sections were dehydrated in 100% ethanol (two changes, 5 minutes each), cleared in xylene (two changes, 5 minutes each), and cover slipped with a permanent mounting medium (DPX). After staining, Nissl bodies within neuronal cytoplasm appeared purple blue, allowing clear visualization of neuronal morphology and cytoarchitecture.

Images were obtained using a Keyence BZ-9000 microscope. Hippocampal volume was measured with the Volumest plugin for ImageJ (NIH) (http://lepo.it.da.ut.ee/~markkom/volumest/). The thickness of the CA1 pyramidal cell layer and dentate gyrus granule cell layer was measured by drawing a straight line perpendicular to the length of the cell layers at specified locations, using ImageJ. Typically, 10 to 12 hippocampal-containing sections were used for each analysis.

### RNA extraction and qPCR

RNA was isolated using TRIzol reagent, followed by further extraction with phenol-chloroform. RNA concentration was determined using a Thermo Nanodrop spectrophotometer. Subsequently, cDNA was synthesized from 500 ng of total RNA using the Maxima RT cDNA synthesis kit. A SYBR Green-containing NEB Luna qPCR mix was used to prepare the qPCR reactions. Quantitative PCR was performed using a Bio-Rad qPCR machine. Primer details are mentioned in **S-Table.5**.

### *In vitro* fibril-induced Tau spreading model

The *in vitro* fibril-induced Tau spreading model was adapted and modified from previously described protocols. Briefly, primary neurons were cultured on coverslips and infected with AAV-P301S PS19 (Virovek, Inc.) on day in vitro 3 (DIV3). Synthetic Tau fibrils (K18/P301L, 100 nM) were added to the culture medium on DIV5. Neurons that did not receive AAV infection (AAV-) and those treated with PBS instead of fibrils served as negative controls. Neurons were incubated with the fibrils for 7 to 10 days. To assess the effect of CST treatment, recombinant CST was added to the culture medium together with the fibrils and replenished once after 3 days. Immunofluorescence staining using the MC1 antibody (1:500) was performed to detect misfolded Tau species. Neurons grown on coverslips were washed with fresh medium, fixed in 4% paraformaldehyde containing 0.1% Triton-X, and processed for standard immunofluorescence staining. CST was added to the culture medium together with the fibrils and replenished once after 3 days. Immunofluorescence staining using the MC1 antibody (1:500) was used to detect misfolded Tau species. Briefly, neurons grown on coverslips were washed with fresh medium, fixed in fresh 4% PFA with 0.1% Triton-X, and then subjected to standard immunofluorescence staining.

### Tau Seeding Assay

HEK 293 cells expressing the human Tau RD P301S FRET biosensor (ATCC CRL-3275) were plated onto Millipore EZ chambered slides at a confluency of 1 × 10^3^ cells per well in complete DMEM (DMEM supplemented with 10% Fetal bovine serum and 100μg/mL penicillin-streptomycin). After 16 hours, the medium was replaced with OptiMEM, and cells were treated with 2 μg of lysate from nTg (n=4), Sal/PS19 (n=4), and CST/PS19 (n=4), prepared in PBS. After 24 hours of treatment, the medium was replaced with complete DMEM. After an additional 24 hours, the medium was removed, and cells were fixed with 4% paraformaldehyde and stained with DAPI. Cells were then mounted and imaged using a Keyence fluorescence microscope with a 40X objective. Four biological replicates per group were included, and four images. Three images from each animal were analyzed using ImageJ for quantification.

### Brain and plasma Pharmacokinetics study of CST

In vivo plasma and brain pharmacokinetics of CST were evaluated in male PS19 mice. Mice received a single intraperitoneal injection of CST at a dose of 5 mg/kg. Blood samples (∼100 μl) were collected via tail vein at 0 (pre-dose), 0.25, 0.5, 1, 2-, 4-, 8-, and 24-hours post-injection. At selected points, CST-treated mice were euthanized, and whole brains were rapidly dissected. One-half of each brain was homogenized in PBS, and the resulting homogenates were used for CST quantification. For sample preparation, 50 μl of plasma or brain homogenate was mixed with 150 μl of methanol, vortexed, and centrifuged to precipitate proteins. The supernatant was removed, and the remaining pellet was reconstituted in 150μl of 10% formic acid in water. Samples were vortexed, centrifuged, and a 100 μl aliquot was collected, diluted 1:1 with 10 mM ammonium bicarbonate, and analyzed by liquid chromatography–tandem mass spectrometry (LC–MS/MS).

### Statistics

In the figure legends, ‘n’ denotes the number of biological replicates for each group. All *in vivo* experiments maintained a 1:1 ratio of male to female subjects. Normality was assessed using the Shapiro-Wilk test in GraphPad Prism version 10.4.1. Data were analyzed using unpaired two-tailed t-tests, one-way ANOVA, or two-way ANOVA. Data are presented as mean values ± SEM.

### Data availability

Original data generated and analyzed during this study are included in this article and are available from the lead corresponding author on reasonable request.

### Code availability

Codes are available at GitHub: sahoo00/snRNASeq-PS19.

## Supporting information

Supplementary_Figures_Legends_BioRxiv_01032026

## Acknowledgments

This work was supported by NIH grants AG080246, AG078635, and AG091126 to SKM; R01-AI155696, R01-GM138385, and UG3TR003355 to DS; R01AG074273 and R01AG078185 to XC, as well as VA RR&D SPiRE grant RX004398 to SKM. SJ was supported by an AFTD Holloway Postdoctoral Fellowship (2022-0002), and DM-M was supported by the AARF-D fellowship from the Alzheimer’s Association. We also acknowledge support from the Wu Tsai Human Performance Alliance (WTHPA) and the Joe and Clara Tsai Foundation. We thank UCSD Shiley-Marcos Alzheimer’s Disease Research Center (ADRC) for providing AD postmortem brain tissues, blood and CSF samples, and Dr. Dennis Dickson for providing CBD postmortem brain tissue samples from the Jacksonville Mayo Clinic Brain Bank. We want to acknowledge ***UCSD*** School of Medicine ***Microscopy Core*** (Grant P30 NS047101).

## Contribution to work

Conceptualization: SKM, SJ

Methodology: SJ, SK, SKM, XC, DM-M, KT, DS

Investigation: SJ, SK, SKM, KT, XC, DM-M, DS

Visualization: SJ, SK, SKM, KT, XC, DM-M, DS

Funding acquisition: SKM, XC, DS, SJ, DM-M

Project administration: SJ, SKM

Supervision: SJ, SKM, XC Writing – original draft: SKM, SJ

Writing-review and editing: XC, SJ, DS, SKM

