## Supplementary_Figures_Legends_BioRxiv_01032026 for "Catestatin ameliorates tauopathy and amyloidogenesis via adrenergic inhibition"

**Supplementary Figures and Legends**

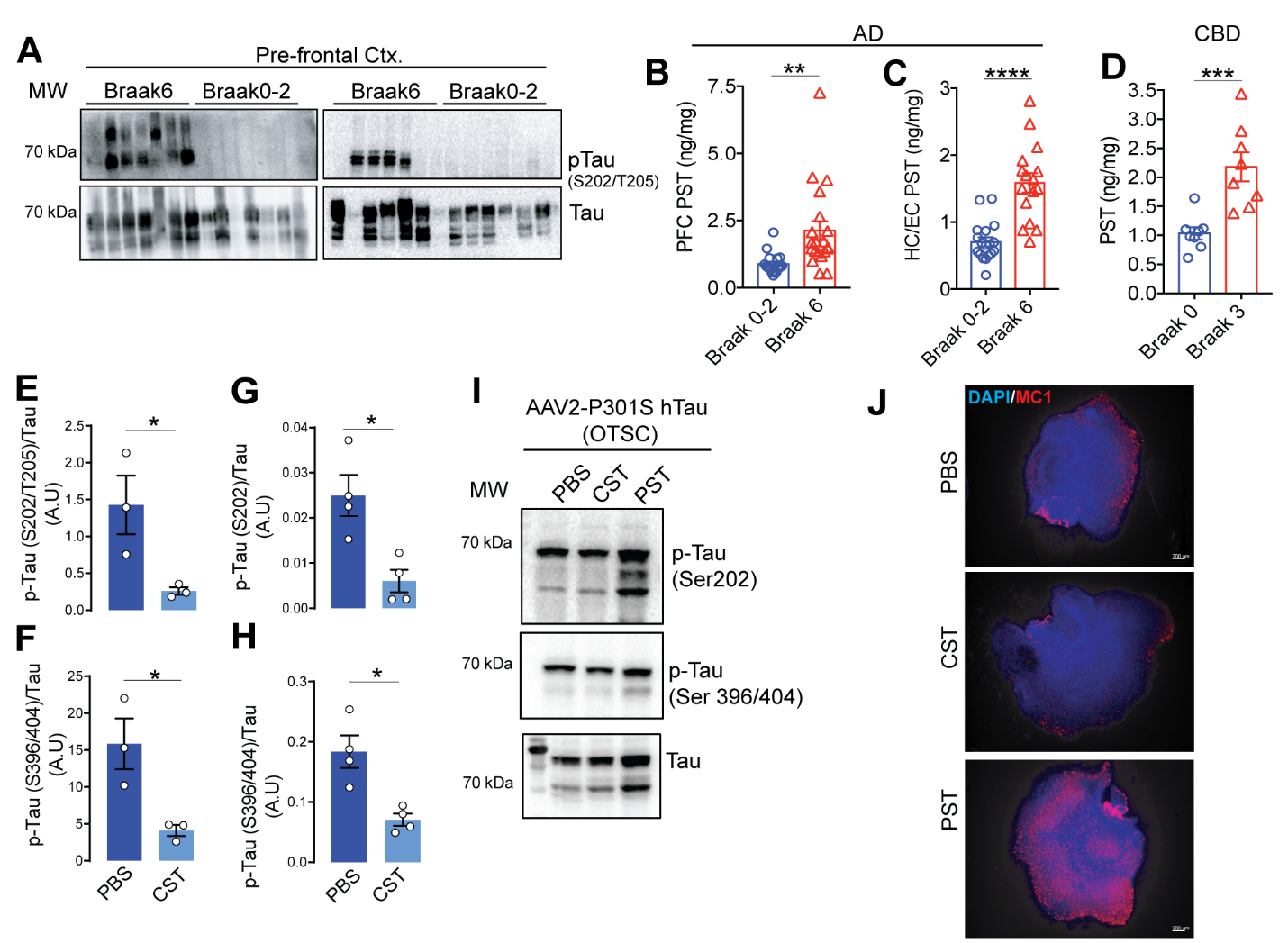

**S-Figure 1. PST levels are increased in Alzheimer’s Disease (AD), CorticoBasal Degeneration (CBD), and PS19 mice, and PST treatment increases Tau phosphorylation. (A)** Representative immunoblots of prefrontal cortex (PFC) samples from AD subjects (Braak 6 vs. Braak 0–2) showing increased p-Tau (S202/T205) in samples with lower PST levels. **(B–D)** PST levels in human brain tissue. (**B**) Quantification of PFC PST (Braak 0-2: n=18, Braak 6: n=20) and (**C**) hippocampal/entorhinal cortex (HC/EC) PST in AD cases shows significantly higher PST levels in Braak 6 than in Braak 0–2 (Braak 0-2: n=18, Braak 6: n=16). (**D**) CBD cases similarly show higher PST in Braak 3 (n=8) relative to Braak 0 (n=8). **(E–H)** CST suppresses Tau phosphorylation in PS19 hippocampal tissue. CST-treated PS19 mice show significantly reduced (**E**) p-Tau at S202/T205 and (**F**) S396/404. (**G&H**) Ratios of p-Tau/total Tau for both epitopes. **(I)** CST reduces and PST increases Tau phosphorylation in the AAV2-P301S hTau (OTSC) model. Western blots show decreased p-Tau (Ser202 and Ser396/404) in hippocampi treated with CST and increased Tau phosphorylation post-PST treatment compared with the PBS control. **(J)** MC1 staining of CST, PST, and PBS-treated OTSC. A.U.: Arbitrary Unit. Data are presented as mean ± SEM; statistical significance determined using appropriate tests *p < 0.05, **p < 0.01, *****p < 0.001, ****p < 0.0001; NS, not significant.

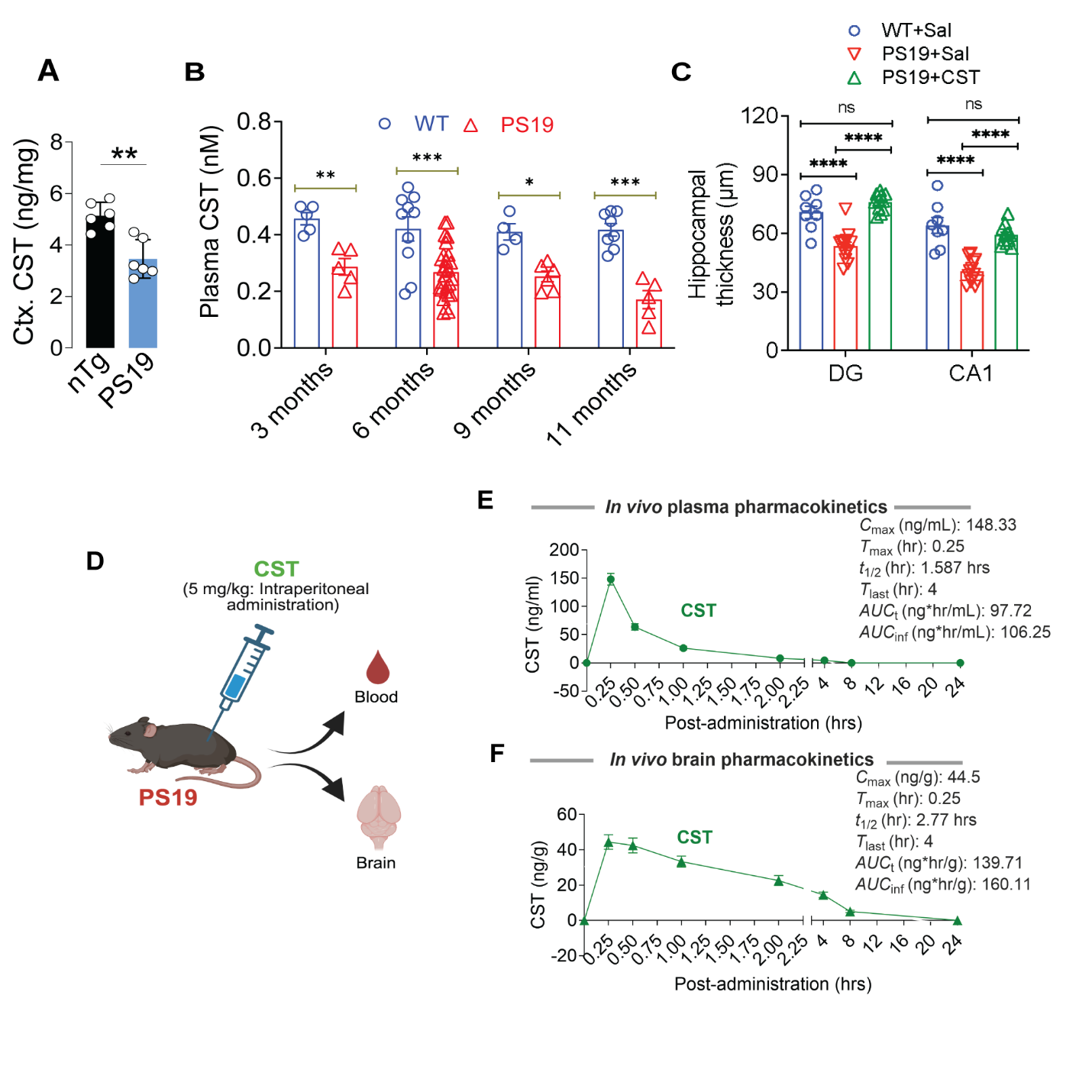

**S-Figure 2. CST levels are decreased in PS19 mice, and CST preserves hippocampal structure in PS19 mice and exhibits rapid plasma uptake with measurable brain penetration.** CST levels are significantly reduced in the cortex (nTg: n=6, PS19: n=6) (**A**) and plasma (**B**) of 9-month-old PS19 mice, indicating that CST deficiency occurs *in vivo* in Tauopathy. (**C**) Sal/PS19 mice show significant thinning of DG and CA1, consistent with Tau-mediated neurodegeneration, whereas CST treatment restores thickness toward nTg levels. **(D–F)** Pharmacokinetics of CST (5 mg/kg, i.p.). (**D**) Schematic of blood and brain collection. (**E**) Plasma PK shows rapid absorption into CST (*T*_max_ = 0.25 hr) and a short systemic half-life (*t*_1/2_ = 1.59 hr). (**F**) Brain PK confirms CNS entry with detectable CST levels and a longer brain half-life (*t*_1/2_ = 2.77 hr), supporting CST’s ability to engage central targets (n=3 at each time point). Data shown as mean ± SEM. Significance indicated as *p < 0.05, **p < 0.01, *****p < 0.001, ****p < 0.0001; NS, not significant.

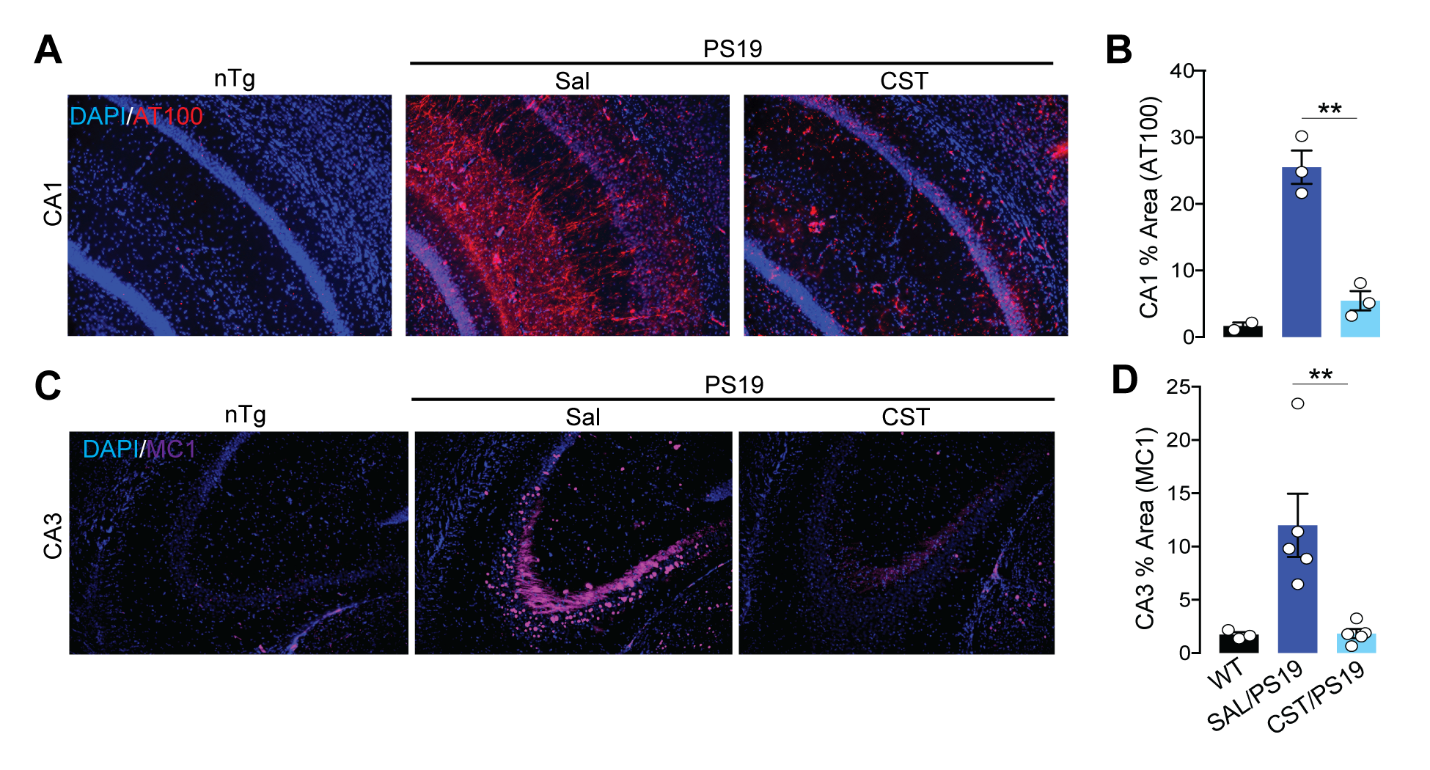

**S-Figure 3. CST reduces hyperphosphorylated and misfolded Tau species in PS19 hippocampus. (A&B)** AT100 immunostaining (p-Tau S212/T214) in CA1. (**A**) PS19 mice show pronounced AT100⁺ Tau pathology compared with nTg controls, reflecting accumulation of late stage hyperphosphorylated Tau. CST treatment markedly decreases AT100 immunoreactivity (nTg: n=2, Sal/PS9: n=3, CST/PS19: n=3). (**B**) Quantification of AT100⁺ area confirms CST-mediated suppression of pathological Tau phosphorylation in CA1. **(C&D)** MC1 immunostaining (conformationally misfolded Tau) in CA3. (**C**) PS19 mice exhibit strong MC1⁺ Tau accumulation, consistent with misfolded Tau species formation. CST reduces MC1 signal intensity and distribution. (**D**) Quantification of MC1⁺ area shows a trend toward decreased misfolded Tau in CST-treated PS19 mice (nTg: n=3, Sal/PS9: n=3, CST/PS19: n=3). Data are shown as mean ± SEM. *p < 0.05, **p < 0.01, *****p < 0.001, ****p < 0.0001; NS, not significant.

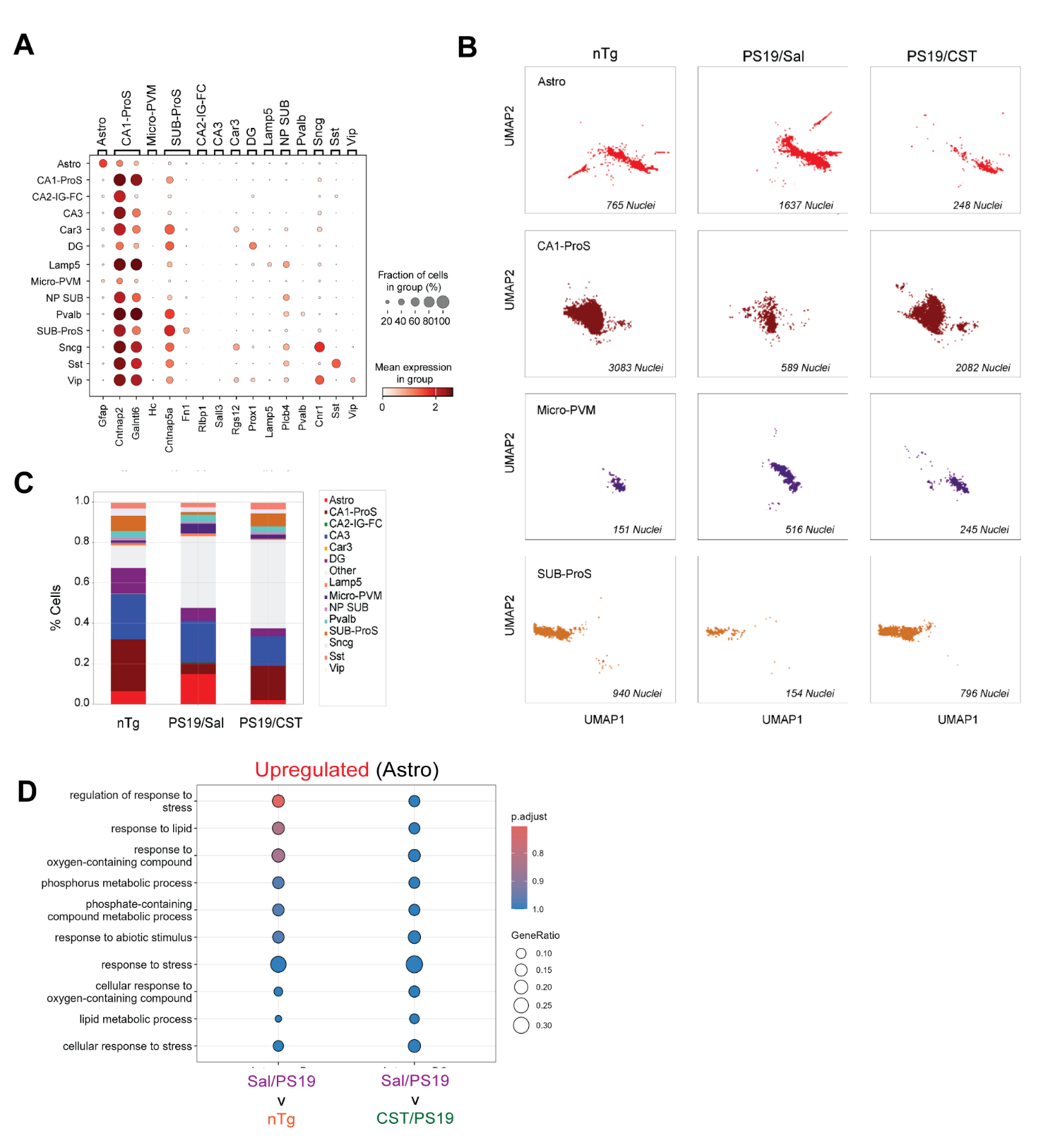

**S-Figure 4. CST restores vulnerable neuronal populations and normalizes transcriptional states in PS19 hippocampus. (A)** Gene signature of the clusters identified and analyzed. **(B)** Cell-type composition derived from snRNA-seq. PS19/Sal mice show reduced proportions of CA1-ProS and SUB-ProS excitatory neurons, with expansion of microglia, reflecting Tau-driven neurodegeneration and inflammatory remodeling. CST treatment reverses these shifts, restoring neuronal populations and reducing microglial abundance toward nTg levels. **(C)** UMAP visualization of key cell classes. PS19/Sal mice exhibit increased astrocyte population, marked loss of CA1-ProS and SUB-ProS nuclei and increased Micro-PVM clustering, whereas CST substantially rescues CA1-ProS and SUB-ProS representation and decreases activated microglia. (**D**) Upregulated pathways in Astro cluster of Sal/PS19 compared to nTg and CST/PS19 indicating an increased metabolic stress in Sal/PS19.

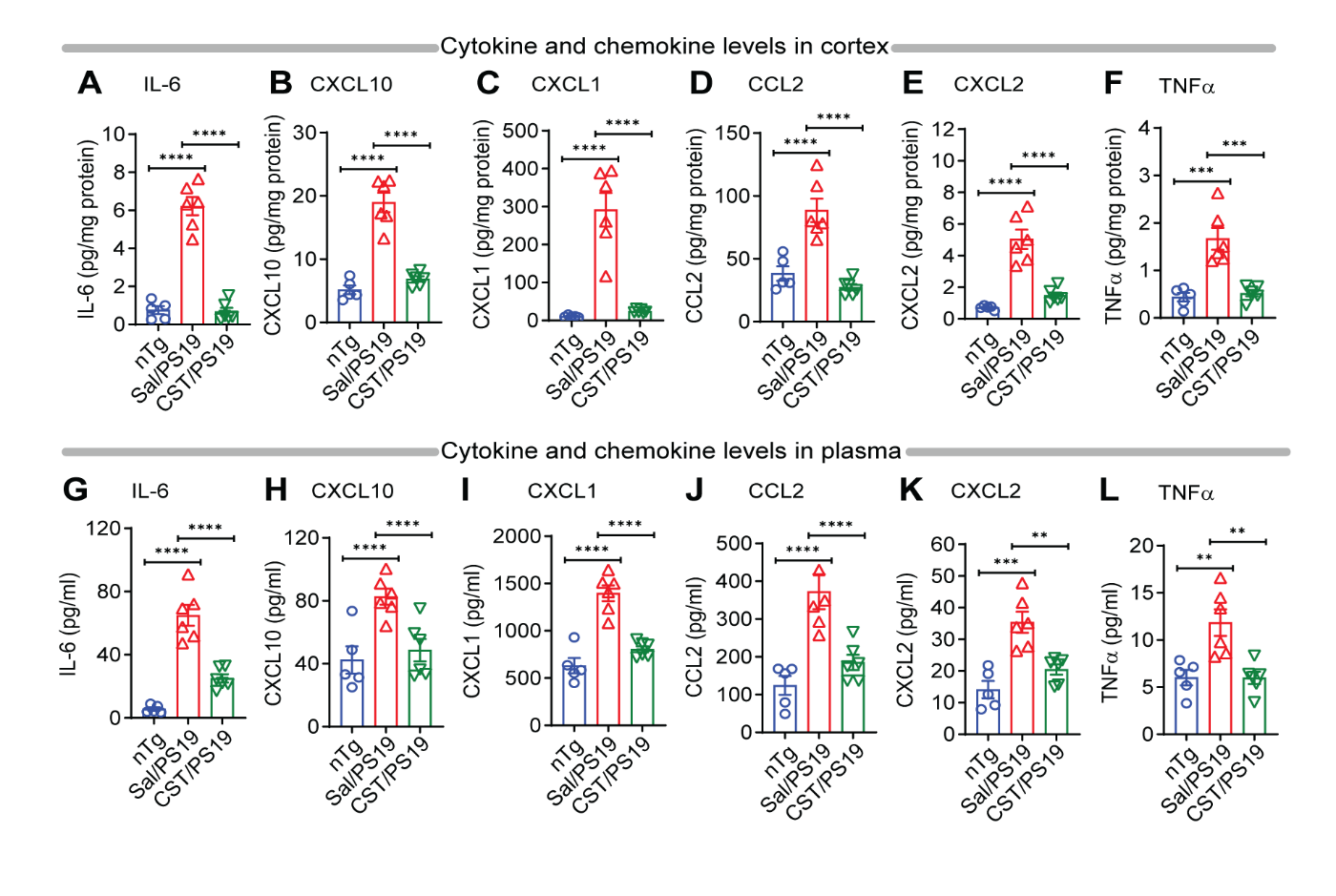

**S-Figure 5. CST suppresses elevations in cortical and systemic levels of inflammatory cytokines/chemokines in PS19 mice. (A–F)** Cortex: PS19 mice show strong upregulation of pro-inflammatory cytokines and chemokines - including (**A**) IL-6, (**B**) CXCL10, (**C**) CXCL1, (**D**) CCL2, (**E**) CXCL2, and (**F**) TNFα - reflecting Tau-driven neuroinflammatory activation. CST treatment significantly reduces all measured factors, indicating potent suppression of central inflammatory signaling (nTg: n=5, Sal/PS19: n=6, CST/PS19: n=6). **(G–L)** Plasma: Systemic inflammation mirrors cortical pathology, with PS19 mice exhibiting elevated circulating (**G**) IL-6, (**H**) CXCL10, (**I**) CXCL1, (**J**) CCL2, (**K**) CXCL2, and (**L**) TNFα. CST markedly lowers these plasma biomarkers, demonstrating that CST attenuates both central and peripheral inflammatory responses (nTg: n=5, Sal/PS19: n=5, CST/PS19: n=6). Data shown as mean ± SEM.

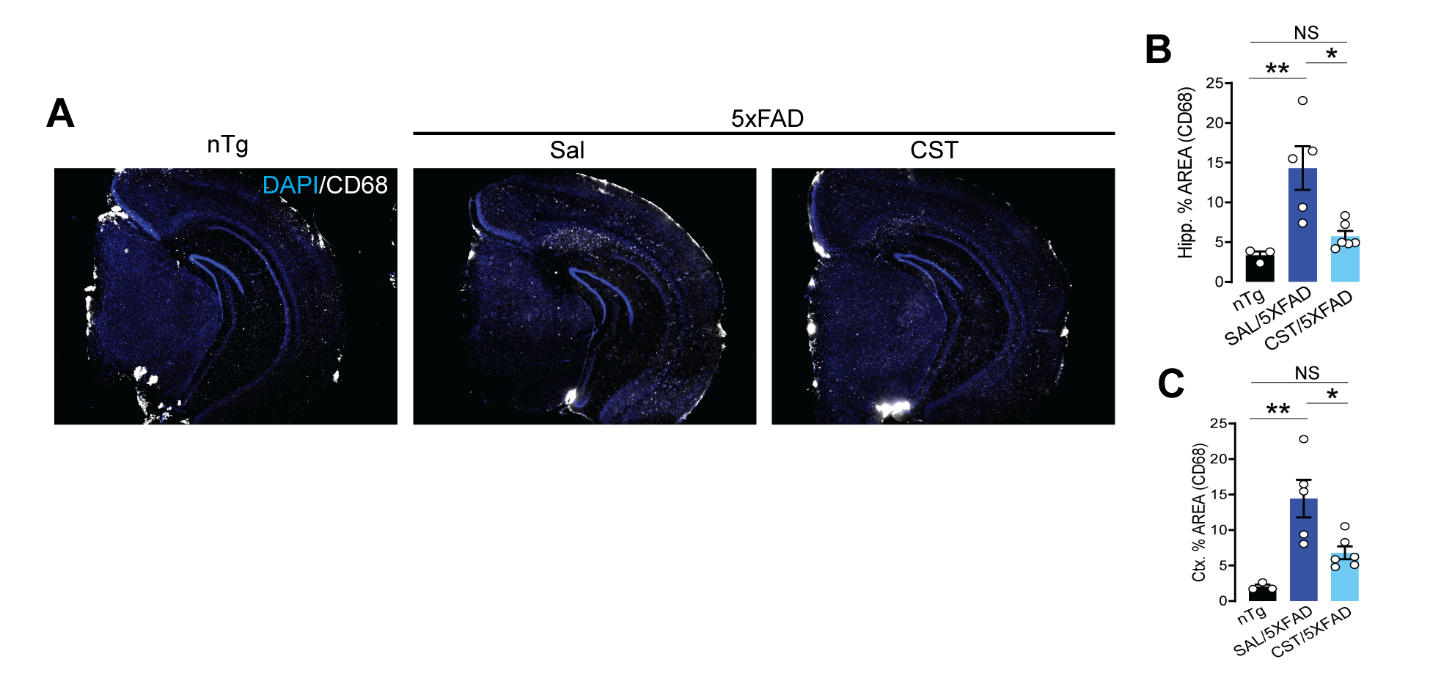

**S-Figure 6. CST reduces microglial activation in the 5xFAD model of amyloid-driven neuroinflammation. (A)** CD68 immunostaining in hippocampal sections from nTg, 5xFAD+Sal, and 5xFAD+CST mice. 5xFAD mice exhibit widespread CD68⁺ microglial activation, reflecting amyloid-induced innate immune activation. CST treatment markedly reduces CD68 signal intensity and distribution. **(B&C)** Quantification of CD68⁺ area in hippocampus (B) and cortex (C). CST significantly lowers microglial activation in both regions, demonstrating its ability to suppress amyloid-associated neuroinflammatory responses (nTg: n=3, Sal/5xFAD: n=5, CST/5xFAD: n=6). Data shown as mean ± SEM. *p < 0.05, **p < 0.01, *****p < 0.001, ****p < 0.0001; NS, not significant.

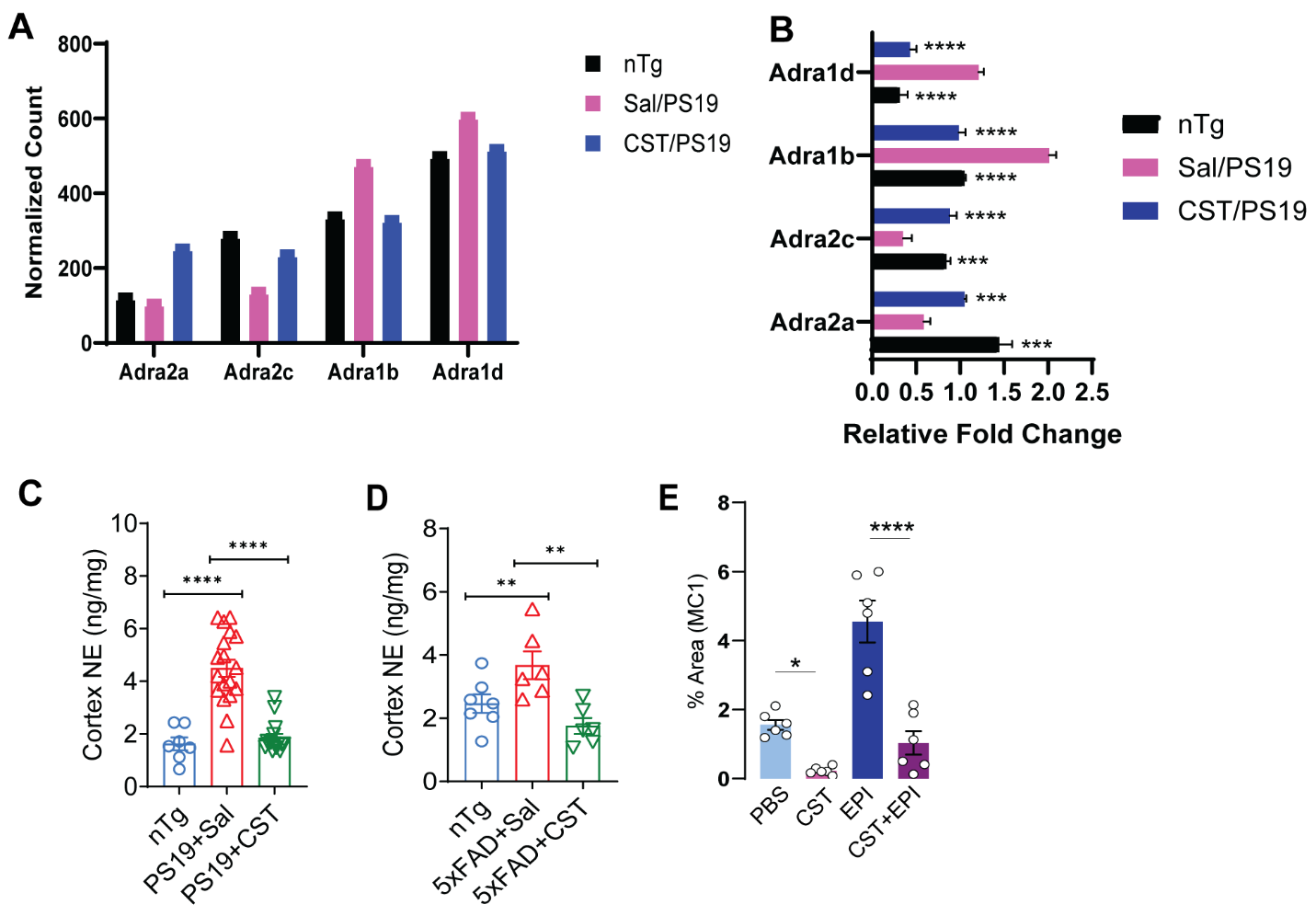

**S-Figure 7. CST counteracts adrenergic stimulation–induced Tau misfolding and integrates with measurements of norepinephrine signaling. (A)** CST treatment reduces ADRA1 and increases ADRA2 expression in PS19 hippocampus. (B) qPCR analysis of ADRA1B, ADRA1D, ADRA2A, ADRA2C in hippocampus of nTg, Sal/PS19, CST/PS19 (n=4). **(C-D)** Cortex Norepinephrine (NE) measurement in PS19 **(C)** and 5xFAD mice **(D)** after CST treatment. **(E)** CST prevents epinephrine (EPI)-induced accumulation of misfolded Tau. Quantification of MC1⁺ Tau in hippocampal sections shows that EPI markedly increases misfolded Tau species, consistent with adrenergic/PKA pathway overactivation. CST alone shows minimal MC1 staining and significantly reduces EPI-driven Tau misfolding when co-administered, indicating that CST antagonizes ADR–mediated pathogenic Tau conformations. Data are shown as mean ± SEM; statistical significance is indicated. *p < 0.05, **p < 0.01, *****p < 0.001, ****p < 0.0001; NS, not significant.
